# A recently formed triploid *Cardamine insueta* inherits leaf vivipary and submergence tolerance traits of parents

**DOI:** 10.1101/2020.06.03.130500

**Authors:** Jianqiang Sun, Rie Shimizu-Inatsugi, Hugo Hofhuis, Kentaro Shimizu, Angela Hay, Kentaro K. Shimizu, Jun Sese

## Abstract

Contemporary speciation provides a unique opportunity to directly observe the traits and environmental responses of a new species. *Cardamine insueta* is an allotriploid species that appeared within the past 150 years in a Swiss village, Urnerboden. In contrast to its two progenitor species, *C. amara* and *C. rivularis* that live in wet and open habitats, respectively, *C. insueta* is found in-between their habitats with temporal water level fluctuation. This triploid species propagates clonally and serves as a triploid bridge to form higher ploidy species. Although niche separation is observed in field studies, the mechanisms underlying the environmental robustness of *C. insueta* are not clear. To characterize responses to a fluctuating environment, we performed a time-course analysis of homeolog gene expression in *C. insueta* in response to submergence treatment. For this purpose, the two parental (*C. amara* and *C. rivularis*) genome sequences were assembled with a reference-guided approach, and homeolog-specific gene expression was quantified by using HomeoRoq software. We found that *C. insueta* and *C. rivularis* initiated vegetative propagation by forming ectopic meristems on leaves, while *C. amara* did not. We examined homeolog-specific gene expression of three species at nine time points during the treatment. The genome-wide expression ratio of homeolog pairs was 2:1 over the time-course, consistent with the ploidy number. By searching the genes with high coefficient of variation of expression over time-course transcriptome data, we found many known key transcriptional factors related to meristem development and formation upregulated in both *C. rivularis* and *rivularis*-homeolog of *C. insueta*, but not in *C. amara*. Moreover, some *amara*-homeologs of these genes were also upregulated in the triploid, suggesting trans-regulation. In turn, Gene Ontology analysis suggested that the expression pattern of submergence tolerant genes in the triploid was inherited from *C. amara*. These results suggest that the triploid *C. insueta* combined advantageous patterns of parental transcriptomes to contribute to its establishment in a new niche along a water-usage gradient.

## 1 Introduction

The molecular basis of speciation has been a central question in biology (Jerry A and H. Allen, 2004). Little is known still about how a new species obtains new traits to adapt to a distinct environment. A major obstacle in studying this is that most speciation events occurred in the past, and thus the traits and the environment at the time of speciation are not directly observable. The difference in traits and environments between current species may represent evolution after speciation rather than the changes that occurred at speciation. A unique opportunity to study speciation in action is contemporary allopolyploid speciation (Soltis and Soltis, 2009; Abbott et al., 2013). During the past 150 years, several cases of polyploid speciation have been documented, for example in *Tragopogon, Senecio, Mimulus, Spartina*, and *Cardamine* (Urbanska et al., 1997; Abbott and Andrew, 2004; Ainouche et al., 2004; Soltis et al., 2004). Because polyploid speciation immediately confers complete or partial reproductive isolation between the new polyploid and progenitor species, a new polyploid species must establish and propagate while surrounded by individuals with different ploidy. To overcome this situation termed “minor cytotype disadvantage”, two traits are suggested to facilitate establishment (Comai, 2005). First, the distinct environmental niche of a polyploid species would reduce competition with progenitor species. Second, clonal vegetative propagation or self-fertilization would assure the persistence of new polyploids at the initial stages because meiotic abnormality is common in newly formed polyploid species. This would be critical for odd-ploidy species including triploids, which often contribute to the formation of higher polyploids via a so-called triploid bridge (Bretagnolle and Thompson J. D., 1995; Ramsey and Schemske, 1998; Mable, 2003; Husband, 2004; Tayalé and Parisod, 2013; Mason and Pires, 2015). Despite the significance of these traits, the underlying molecular mechanisms are yet to be studied.

The contemporary polyploid *C. insueta* belongs to the genus *Cardamine*, which has long been studied for ecological polyploid speciation (Howard, 1948; Hussein, 1948), and represents adaptive radiation by recurrent polyploidization along water-usage gradients (Shimizu-Inatsugi et al., 2016; Akiyama et al., 2019). A major advantage to studying *Cardamine* is that it is closely related to the model plant *Arabidopsis thaliana*, and a reference genome assembly of *Cardamine hirsuta* (Gan et al., 2016) is publicly available, thus functional and genomic data of these model species are readily available. One allotriploid species in *Cardamine, C. insueta* (2*n* = 3*x* = 24; RRA), is a textbook example of contemporary speciation discovered by Urbanska and Landolt in 1974 (Urbanska-Worytkiewicz and Landolt, 1974b). It was formed by the hybridization of two progenitor diploids *Cardamine amara* (2*n* = 2*x* = 16; AA) and *Cardamine rivularis* (2*n* = 2*x* = 16; RR, belonging to *C. pratensis* complex sensu lato) approximately 100–150 years ago at the valley of Urnerboden in the Swiss Alps (Urbaska-Worytkiewicz and Landolt, 1972; Urbanska-Worytkiewicz and Landolt, 1974b; Urbanska et al., 1997; Mandáková et al., 2013; Zozomová-Lihová et al., 2014) (**Fig. S1*A***). The two diploid progenitors have distinct ecological habitats. While *C. amara* grows in and beside water streams, *C. rivularis* inhabits slightly moist sites, avoiding permeable and fast drying soil (Urbanska-Worytkiewicz and Landolt, 1974b, 1974a) (**Fig. S1*B***). Around the end of the 19th to the early 20th centuries, the deforestation and land-use conversion to grazing induced the hybridization of these two diploids to produce the triploid species *C. insueta*, which is abundant in manured hay-meadows (Urbaska-Worytkiewicz and Landolt, 1972; Urbanska et al., 1997; Mandáková et al., 2013). Cytogenetic studies suggested that *Cardamine insueta* served as a triploid bridge in the formation of pentaploid and hexaploid *C. schulzii* by the further hybridization with autotetraploid *C. pratensis* (sensu stricto, 2*n* = 2*x* = 30; PPPP; hypotetraploid derived from a chromosomal fusion) in Urnerboden (Mandáková et al., 2013).

The propagation of triploids mainly depends on vegetative propagation for two reasons, high male sterility *per se* and hay cutting and grazing in flowering season (Urbanska et al., 1997). One of the progenitor species, *C. rivularis*, can produce plantlets on the surface of leaves and nodes by ectopic meristem formation, which is a common feature of the *C. pratensis* complex (Smith, 1825; Salisbury, 1965; Dickinson, 1978). This characteristic is inherited by *C. insueta*, enabling it to be a dominant species at the site despite its ploidy level (Urbanska-Worytkiewicz and Landolt, 1974a; Urbanska et al., 1997). This type of leaf vivipary is only found in a limited number of angiosperms and assumed to contribute to population establishment in polyploids (Dickinson, 1978). In this sense, the trait of leaf vivipary can be considered a key factor for the establishment of this triploid.

Another interesting aspect of *C. insueta* establishment is its ecological niche shift relative to its progenitor species. Genus *Cardamine* is known to include many submergence tolerant species including *C. amara* (Shimizu-Inatsugi et al., 2016; Akiyama et al., 2019). An allotetraploid *C. flexuosa*, derived from *C. amara* and *C. hirsuta* diploid progenitors, was shown to inherit parental traits and be successful in a wider soil moisture range (Shimizu-Inatsugi et al., 2016; Akiyama et al., 2019). The transcriptomic response of *C. flexuosa* to submergence or drought stress was shown to be combined although attenuated compared to its progenitor species, which could confer the wider tolerance found in the polyploid. Even though the niche separation between *C. rivularis* and *C. insueta* is not yet clearly illustrated, our field observations are consistent with this hypothesis.

In this study, we focused on the time-course gene expression pattern of the triploid *C. insueta* and its two diploid progenitors during submergence treatment, which induces both water stress and ectopic meristem formation on leaves. To study the time-course data of homeologs, we employed bioinformatic methods of variably expressed genes because data points of a time-course are not independent and serve partly as replicates (Yamaguchi et al., 2008; Shin et al., 2014). Here we combined the time course analysis with subgenome-classification bioinformatic workflow of HomeoRoq (Akama et al., 2014), and detected variably expressed homeologs (VEH) during the treatment. We address the following specific questions:

1. What is the expression rate and the ratio of homeologous genes in triploid species in response to submergence, either genome-wide or between each homeologous gene pair?
2. Which gene ontology categories are enriched in VEH? Do they reflect the phenotypic trait of each progenitor species or the triploid? How does *C. insueta* combine the expression patterns of the two progenitors?

## 2 Materials and Methods

### 2.1 Plant materials and RNA sequencing

*Cardamine insueta*, *C. amara*, and *C. rivularis* plants used in this study were collected from Urnerboden. All plants were grown together in a plant cultivation room with 16 hr light and 8 hr dark cycle. The plants were planted in single pots, placed on trays, and watered from below.

Submergence treatment was started in the morning at 07:00. Two mature leaves were detached and submerged in water. We isolated RNA from the floating leaflets of the three species at nine time points after the start of submergence treatment (0 hr, 2 hr, 4 hr, 8 hr, 12 hr, 24 hr, 48 hr, 72 hr, and 96 hr) using Qiagen RNeasy kit (Qiagen, Maryland, U.S.A.). RNA quality was assessed by Bioanalyser Nanochip (Agilent, Santa Clara, U.S.A.) and libraries quantified by Qubit (ThermoFisher, Waltham, U.S.A.). In total 27 libraries (3 species x 9 time points) were prepared according to NEBNext Ultra™ Directional RNA Library Prep Kit for Illumina (New England Biolabs, Ipswich, U.S.A.) followed by paired end sequencing (100bp x 2) on a HiSeq2000 with a HiSeq Paired-End Cluster Generation Kit and HiSeq Sequencing Kit (Illumina, San Diego, U.S.A.). Trimmomatic (ver. 0.36) (Bolger et al., 2014) was used for discarding the low-quality reads with parameters of “PE -threads 4 -phred33 ILLUMINACLIP:adapters.fa:2:30:10 LEADING:20 TRAILING:20 SLIDINGWINDOW:4:20 MINLEN:50”.

### 2.2 Reference sequence assembly

The reference sequences of A-genome and R-genome were assembled by SNP substitution at coding regions from the *C. hirsuta* genome (Gan et al., 2016) with the following steps. To assemble the reference sequence of A-genome, first, we pooled all RNA-Seq reads of the nine RNA-Seq samples of *C. amara*. Second, we mapped the reads onto the reference sequence (i.e., H-genome) using STAR (ver. 2.3.0e) (Dobin et al., 2013). Third, we detected single-nucleotide polymorphisms (SNPs) and short indels from the mapping result using samtools (ver. 0.1.18) (Li et al., 2009). SNPs and indels. were defined as the polymorphic loci where at least 80% of reads have the alternative nucleotides. Fourth, we replaced the nucleotides on the reference with the alternative nucleotides, if the alternative nucleotide was covered by at least five reads. Finally, the gene annotations of the assembled sequence were converted from the H-genome annotations with the replacement information. To improve the accuracy of sequence, we used the assembled sequence as a reference sequence, and repeated steps two through five, nine times. The resulting A-genome was used for the mapping of individual RNA-seq data from all three species. The R-genome was also reconstructed with the same protocol. As a result, 1,496,561 and 1,484,186 SNP regions on the H-genome were replaced for *C. amara* and *C. rivularis*, respectively.

### 2.3 Evaluation of HomeoRoq classification confidence using diploids

We used HomeoRoq (ver. 2.1) (Akama et al., 2014) to classify genomic origins of homeolog-specific reads in the nine *C. amara* and *C. rivularis* samples. Following the HomeoRoq pipeline, for each *C. amara* sample, we used STAR to map reads onto the A-genome and R-genome and used HomeoRoq to classify reads as *A-origin*, *R-origin*, and *unclassified*. Then, we calculated the percentage of misclassified reads (i.e., the reads that were classified as *R-origin*). Similarly, we used HomeoRoq to calculate the percentage of misclassified reads (i.e., the reads that were classified as *A-origin*) in each *C. rivularis* sample.

### 2.4 Homeolog expression quantification and A-origin ratio definition of triploid

We used HomeoRoq to analyze the nine *C. insueta* samples. For each *C. insueta* sample, we used STAR to map reads onto A-genome and R-genome and used HomeoRoq to classify reads as *A-origin*, *R-origin*, and *unclassified*. Then, we customized HTSeq (Planet et al., 2012) to count the number of read pairs that mapped on homeolog region for *A-origin*, *R-origin*, and *unclassified* reads of each *C. insueta* sample separately. In the customized HTSeq, if a read mapped on the region overlapped by multiple homeologs, a read was divided by the number of homeologs.

To calculate the number of fragments per kilobase mapped (FPKM) for *C. insueta* samples (I_A_ and I_R_ samples), we first allocated the *unclassified* reads into *A-origin* and *R-origin* reads with A-origin ratio. A-origin ratio of homeolog *h* at the time point *s* was defined as 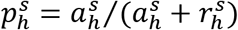, where 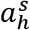 and 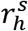 are the numbers of *A-origin* and *R-origin* reads of homeolog *h* at the time point *s*, respectively. Thus, the number of *A-origin* reads after *unclassified* reads allocation 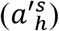 was calculated as 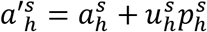, where 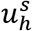 is the number of *unclassified* reads of homeolog *h* in sample *s*. Similarly, 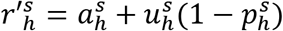 for *R-origin* reads. Then, FPKM of *A-origin* reads of homeolog *h* in sample *s* was calculated as 10^9^ 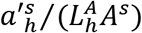, where 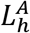 is the length of homeolog *h* on A-genome and *A^s^* is the total number of *A-origin* reads in sample *s*; likewise, FPKM of *R-origin* reads was calculated as 10^9^ 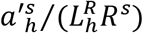, where 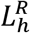 is the length of homeolog *h* on R-genome and *R^s^* is the total number of *R-origin* reads in sample *s*.

In addition, FPKM of progenitors were calculated from the total number of reads (i.e., 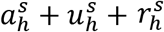). Therefore, FPKM of *C. amara* and *C. rivularis* were calculated as 10^9^ 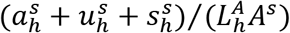 and 10^9^ 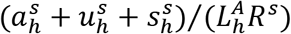, respectively.

### 2.5 Expressed homeologs and PCA analysis

An expressed homeolog was defined as a homeolog with FPKM > 1.0. A homeolog expressed in a sample (i.e., either *amara*-derived in *C. insueta* (I_A_), *rivularis*-derived in *C. rivularis* (I_R_), *C. amara* or *C. rivularis*) was defined as a homeolog with FPKM > 1.0 at least at one of the nine time points. In total, 21,131 homeologs were expressed at least in one sample. PCA was performed against log_10_-transformed FPKM of these 21,131 expressed homeologs. To avoid calculating log_10_0, the log_10_-transfromed FPKM was truly calculated as log_10_(FPKM + 1).

### 2.6 Identification of variably expressed homeologs (VEH) and Gene ontology (GO) enrichment analysis

Mean and coefficient of variation (CV) were calculated from log_10_-transformed FPKM over the nine time points. VHE was defined as an homeolog satisfied the mean > 1.0 and the CV > 0.20. We identified from I_A_, I_R_, *C. amara*, and *C. rivularis* samples, separately.

Gene ontology (GO) enrichment analysis was performed for the four variably expressed homeolog (VEH) sets with R packages clusterProfiler (ver. 3.12.0) and org.At.tair.db (ver. 3.8.2) (Yu et al., 2012). To remove redundancies of GO categories, only GO categories which are associated with 10– 500 *Cardamine* homeologs and below the third level in the GO category hierarchy were used. The threshold FDR = 0.1 was used for cutoff of significantly enriched GO categories.

## 3 Results

### 3.1 Plantlet induction on *C. insueta* and *C. rivularis* leaves by submergence

At the field of Urnerboden valley, we could scarcely observe normal seed setting on *C. insueta*, but small plantlets on leaves were frequently observed after flowering, as described previously (Urbanska, 1977). We also observed small plantlets on the leaves of *C. rivularis*. In contrast, *C. amara* does not form plantlets on leaves, rather adventitious roots and shoots were formed from rhizomes. In the natural habitat, the plantlet formation of *C. rivularis* and *C. insueta* can be seen at flowering to post-flowering season (Salisbury, 1965; Urbanska, 1977). It was also reported that *C. pratensis* (which is closely related to *C. insueta* or considered the same species) tend to bear more plantlets on the leaves in damper sites than in drier sites (Salisbury, 1965), implying that high moisture could be the trigger for meristem formation. Thus, we tested plantlet induction by submergence treatment using dissected leaves with this trio of species in the lab. We detached mature leaves from mother plants propagated in a climate chamber and floated the leaves on water. Within 16 h, we observed the activation of dormant shoot meristems and initiation of ectopic root meristems, which formed visible plantlets on *C. rivularis* leaves 96 hours after submergence (**Fig. S2, Dataset S1**). Induction of ectopic plantlets followed a similar time-course in *C. insueta*. In contrast, plantlet induction was not observed on the leaves of *C. amara*. In addition, during the 96-hr treatment, no symptoms of necrosis appeared on any of the leaves, suggesting that all three species have some submergence tolerance for at least 96 hr.

### 3.2 Gene annotation on the two diploid progenitor reference sequences

To detect how homeologous genes are expressed in plantlet induction and submergence treatment, we harvested time-course RNA-Seq samples of *C. insueta* and diploid progenitor leaves at nine time points after initial submergence (i.e., 0, 2, 4, 8, 12, 24, 48, 72, and 96 hr) (**Fig. S3*A***). We harvested the first lateral leaflet pair in young leaves with no ectopic plantlets. To quantify homeolog-specific gene expression, we assembled the genomes of *C. amara* (A-genome) and *C. rivularis* (R-genome), respectively, using the same pipeline of a reference-guided approach using RNA-Seq reads (**Fig. S3*B***). The genome sequence of a close relative, *C. hirsuta* (H-genome) (Gan et al., 2016), was used as a reference. The A-genome structure is reported to be almost perfectly collinear with that of H-genome, except for one pericentric inversion at chromosome 1, by cytological studies (Mandáková et al., 2013, 2014). The genome structures of the A-genome and R-genome are also similar to each other (Mandáková et al., 2013). The length of assembled reference sequences of A-genome and R-genome are 198,651,635 and 198,654,862 nucleotides, respectively, which are nearly the same as the length of the original H-genome (198,654,690 nucleotides). We also annotated the orthologous genes of *C. amara* and *C. rivularis* according to the information of *C. hirsuta* H-genome. In total, we found 23,995 and 24,115 genes covered by at least one read among the nine time points on the assembled A-genome and R-genome, respectively. These gene sets, which correspond to 81.5% and 81.7% of 29,458 genes in H-genome, respectively, were defined as expressed and used for the following analysis.

### 3.3 Expression ratio from each subgenome is consistent with the number of chromosomes

We applied the HomeoRoq analysis pipeline (Akama et al., 2014) to classify the origin of each RNA-seq read of *C. insueta* samples to either *A-origin* (i.e., the genomic origin of the read is A-subgenome) or *R-origin* (**Fig. S3*C***). After filtering for read quality, 10.6 million read pairs on average among the nine samples could be classified as homeolog-specific read pairs (**Dataset S2**). Of the total homeolog-specific read pairs in the *C. insueta* 0 hr sample, 27.3% and 56.7% of read pairs were classified as *A-origin* and *R-origin*, respectively. The remaining 16.0% of read pairs could be classified to neither *A-origin* nor *R-origin* (*unclassified*) due to the lack of SNPs or the identical sequence on the correspondence region. As a whole genome, the ratio of *A-origin* to *R-origin* reads was approximately 1:2. When we analyzed all samples from the other eight time points, we observed a slight increase in the proportion of *A*-origin reads in correlation with the time point, from 1:2.07 at 0 hr to 1:1.90 at 96 hr (**Dataset S2**). Instead of this minor transition, the expression ratio between subgenomes remained A:R ≈ 1:2 with *C. insueta* samples at all time points, indicating that the expression ratio from each subgenome is consistent with the number of chromosome regardless of the submergence treatment.

### 3.4 Most homeolog pairs were expressed in proportion to the subgenomes in *C. insueta*

To investigate the proportion of expression levels of homeolog pairs in *C. insueta*, we quantified the expression level of each homeolog pair at each time point. We found that (i) the correlation between the expression levels of homeolog pairs was higher than 0.81 at any time point (**Fig. 1*A* and Fig. S4**). However, (ii) the expression levels of most homeologs expressed from the A-subgenome (A-homeolog) were approximately half that of R-homeologs. To understand the proportion of expression levels of homeolog pairs in detail, we calculated A-origin ratio—the proportion of A-homeolog expression level to the total A-homeolog and R-homeolog expression levels—for all homeolog pairs at each time point. We found that the distribution of A-origin ratios had a gentle peak at the position of 0.33 at all time points (**Fig. 1*B* and Fig. S5**). This result suggests the expression ratio of the majority of homeolog pairs is consistent with the copy number, i.e. the subgenome-set numbers of the triploid. In addition, we found two sharp peaks at both edges, the positions of 0.0 and 1.0, of A-origin ratio, which represent the homeologs only expressed in of either subgenome.

**Figure 1.**
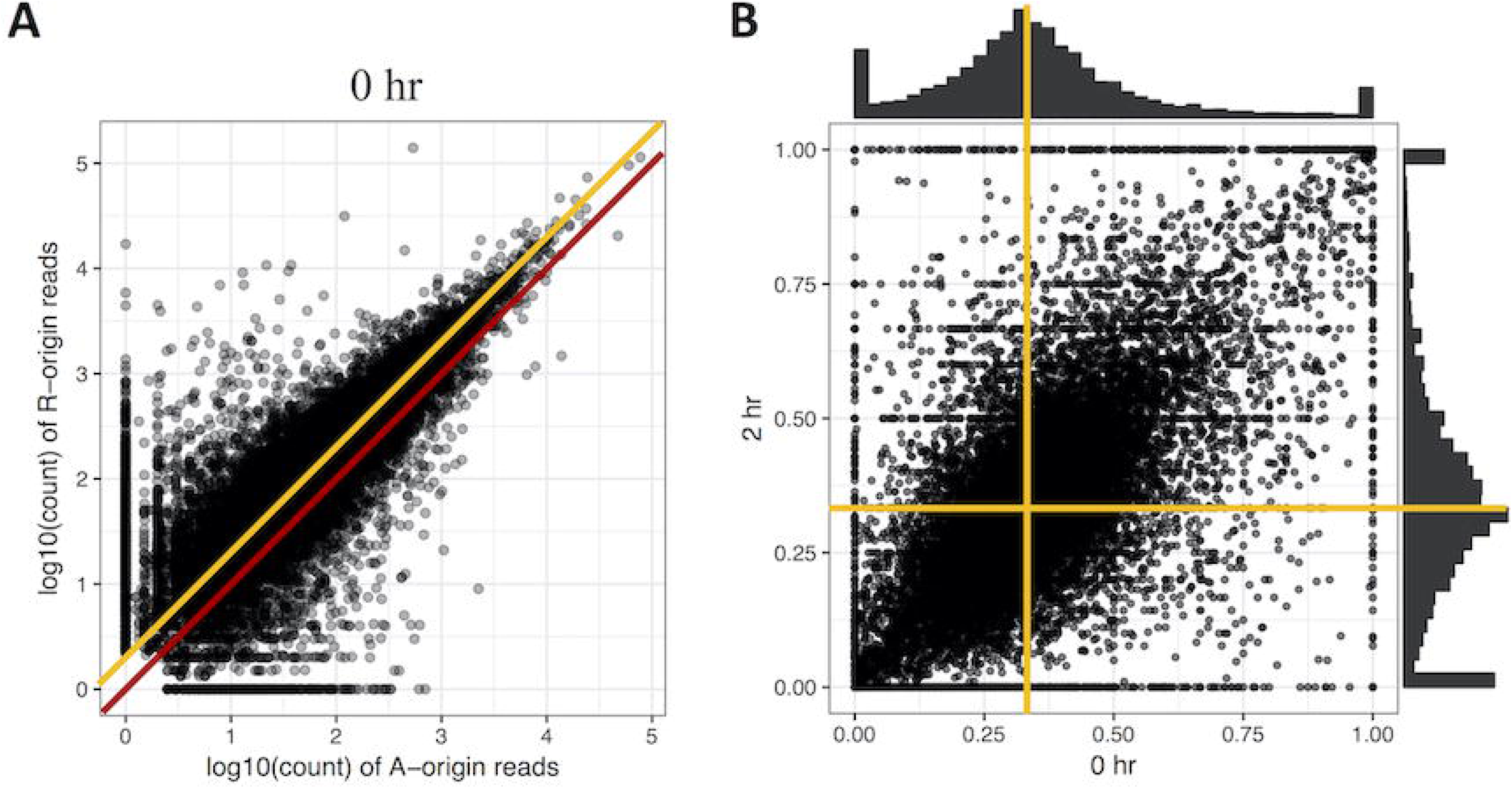
Comparison of homeolog expression from A- and R-subgenomes. **A** Expression ratio between A- and R-homeologs before submergence treatment in the triploid *C. insueta*. Each dot shows the relation between the log10-transformed A-origin and R-origin read of a homeolog pair at 0hr point. Only the homeolog pairs with FPKM > 1.0 in either I_A_ or I_R_ samples are shown. The red line represents the ratio A:R=1:1, and the orange line represents the ratio A:R=1:2. **B** Comparison of A-origin ratios between two time points, 0 hr and 2 hr, in the triploid *C. insueta*. Each point shows the A-origin ratios of a homeolog pair at 0 hr and 2 hr. The orange lines represent the position of A-origin=0.33 at each time point.

Additionally, to investigate whether the A-origin ratio changes during the submergence treatment, we compared the A-origin ratio distributions between different time points. The patterns of all time points were correlated to each other, with the least coefficiency (0.66) between 0 hr and 2 hr (**Fig. 1*B* and Table S1**). This result indicates that A-origin ratios did not change drastically in most homeolog pairs by the submergence treatment, but a limited number of homeolog pairs change the expression balance.

### 3.5 The whole genome expression pattern of each *C. insueta* subgenome is closer to that of its progenitor genome

To gain an overview of how homeologous gene expression varies at the whole genome level among *C. insueta* and the progenitor species *C. amara* and *C. rivularis*, we conducted principal component analysis (PCA). PCA was performed against the log_10_-transformed FPKM of 21,131 expressed homeologs (**Fig. 2**). We found that the first principal component (PC1) grouped samples into two groups: the one with A-homeologs of *C. insueta* (*I_A_*) and *C. amara* (*A*) samples and the other with R-homeologs of *C. insueta* (*I_R_*) and *C. rivularis* (*R*) samples. In addition, we also found that the second principal component (PC2) grouped samples into two groups: one consisting of polyploid samples (*I_A_* and *I_R_* samples, lower side of **Fig. 2A**) and the other consisting of diploid samples (*A* and *R* samples, upper side of **Fig. 2A**). By PC1 and PC2, the samples were grouped into four clusters according to the subgenome type. In contrast, by PC2 and the third principal component (PC3), we observed the transition according to the treatment time, showing a characteristic transition from 0 to 12 hr, and the recurrence of 24, 48, 72 and 96 hr samples towards 0 hr samples in each subgenome, which might reflect the combined effect of submergence stress and circadian rhythm (**Fig. 2B**). The result of PC1 suggests that the majority of the homeologs of *I_A_* and *I_R_* should retain a similar expression pattern to each parent, *A* and *R*. When we focus on PC2, the distance between *R* and *I_R_* is slightly closer than that between *A* and *I_A_*. This might reflect the difference in the number of subgenome sets in the triploid, *A*:*R*=1:2, implying a stronger effect from the pattern with more subgenome sets.

**Figure 2.**
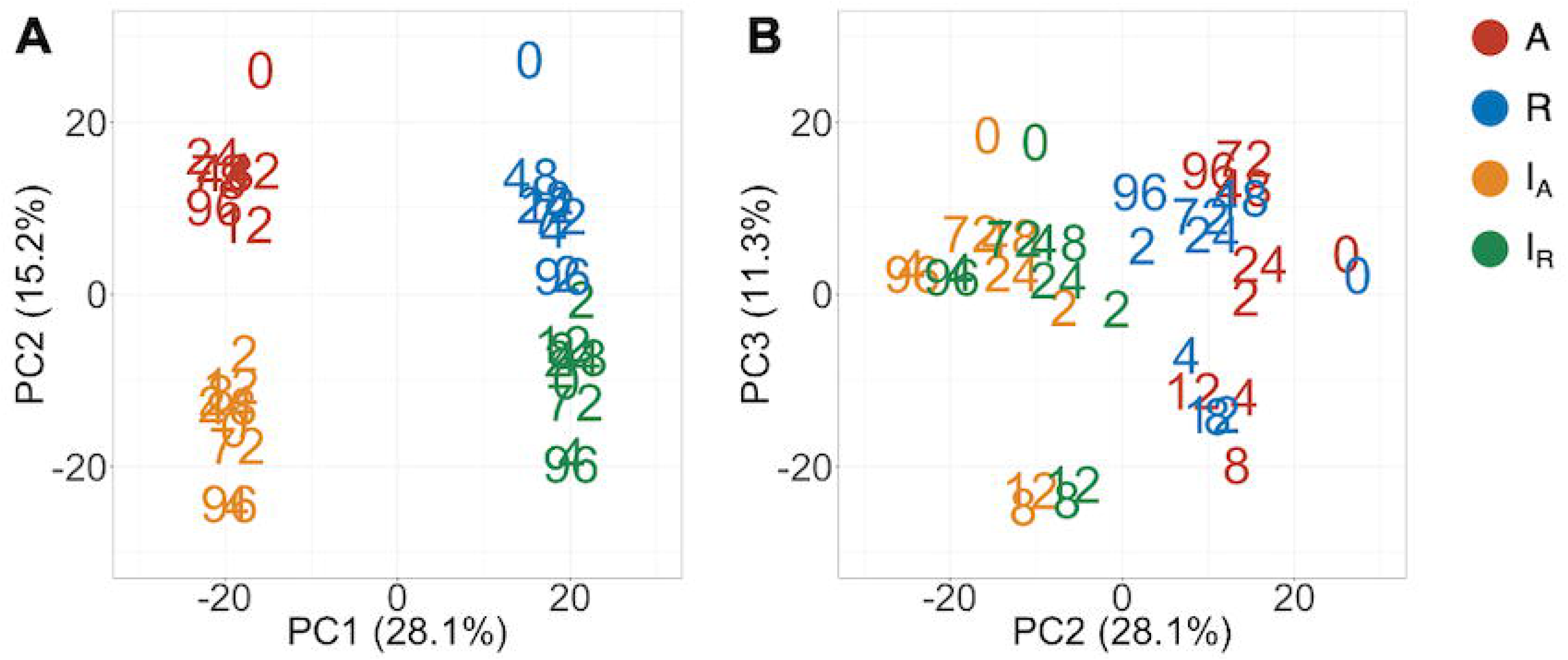
Principal component analysis of the expressed homeologs/genes in *C. insueta* (I_A_), *C. insueta* (I_R_), *C. amara* (A) and *C. rivularis* (R) samples at 9 time points. PCA was performed against log10-transformed FPKM of 21,131 expressed homeologs. The two plots show the relation between PC1-PC2 (**A**) and PC2-PC3 (**B**). The colors represent genome/subgenome, and the numbers represent the time points after the start of submergence treatment.

### 3.6 VEHs related to submergence and their GO enrichment analysis

To understand the difference among species in plantlet formation on the leaf and in submergence response, we focused on the homeologs with a higher expression change during the treatment. Standard tools to identify differentially expressed genes between different conditions are not directly applicable to time-course data, in which expression levels of neighboring time points may be highly correlated. We defined variably expressed homeologs (VEHs) according to the coefficient of variation (CV) among the expression levels of the nine time points, since CV is used for identifying variably expressed genes in various studies involving time-course analysis (Czechowski et al., 2005; Yamaguchi et al., 2008; Shin et al., 2014; Zhao et al., 2017). We identified 1,194, 1,144, 1,030, and 1,063 VEHs from I_A_, I_R_, A and R genome/subgenome with the cutoff CV > 0.2 throughout the treatment, respectively (**Dataset S3**). We visualized the patterns by focusing on two genes that were expected to be affected (**Fig. S6**). The genes associated with ethylene-response such as *ERF1* (AT3G23240) (Chao et al., 1997; Solano et al., 1998) and circadian rhythm such as *CCA1* (AT2G46830) (Alabadí et al., 2001) were identified as VEHs in all samples, which should reflect the ethylene-response to submergence and circadian rhythm response, respectively. The expression pattern of these two homeologs were similar among all four VEH sets from *I_A_*, *I_R_*, *A* and *R* (**Fig. S6**). In addition to these common VEHs, we also found more homeologs identified as VEHs only in one to three samples (**Fig. S7**).

To investigate the biological processes of VEH sets of *I_A_*, *I_R_*, *A* and *R*, we performed gene ontology (GO) enrichment analysis against the four VEH sets (**Table 1**, **Dataset S4**). The numbers of enriched GO categories were 146, 155, 160, and 181, respectively for *I_A_*, *I_R_*, *A* and *R*. A value of negative log10(*q*-value) more than 1 was defined as significant, and a higher value indicates stronger enrichment. We found that some of the GO categories related to water stress, including GO:0006066 (alcohol metabolic process), GO:0009723 (response to ethylene) and GO:0009414 (response to water deprivation), were enriched in all four VEH sets (**Table 1**). As gas diffusion rates are restricted under water, submergence of plants induces ethylene accumulation and low oxygen availability, which could result in the reorganization of the ethylene-response pathway and fermentation pathway (e.g. anaerobic respiration and alcohol metabolism). The enriched categories GO:0006066 and GO:0009723 indicate that *I_A_*, *I_R_*, *A*, and *R* all respond to ethylene and hypoxia signals with the submergence treatment. Two alcohol related categories (GO:0006066 alcohol metabolic process and GO:0046165 alcohol biosynthetic process) were more strongly enriched in *A* and *I_A_*, which was two orders of magnitude higher than I_R_ and R (>2 difference in negative log10(*q*-value) in **Table 1**). In addition, GO:0009414 (response to water deprivation), which encompasses the expression changes of aquaporin genes and ethylene-responsive genes (**Dataset S5**), was enriched.

**Table 1.**
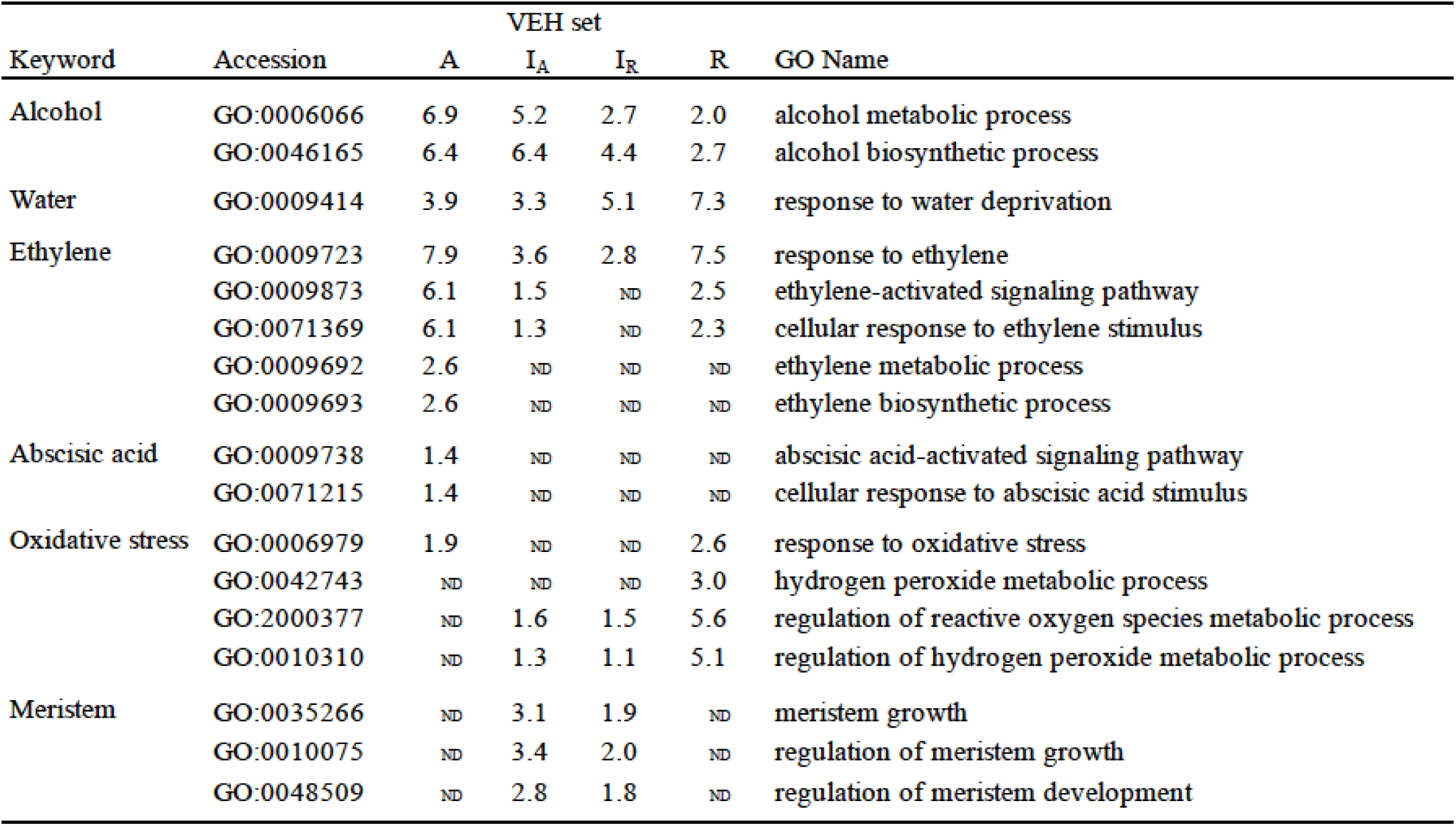
The negative log10(q-value) of the enriched GO categories in each VEH set described in the manuscript. This data is the extract from the list of all enriched GOs (Dataset S4). ND means that the category was not detected as enriched by the threshold FDR = 0.1.

In contrast, some GO categories related to submergence stress were only above the significance threshold in part of the four VEH sets with various combinations (**Table 1**). The two categories related to ethylene metabolism, GO:0009873 (ethylene-activated signaling pathway) and GO:0071369 (cellular response to ethylene stimulus), were not detected in I_R_ but all other three. All these ethylene related GO categories were most strongly enriched in *A*, suggesting larger number of genes are detected than other VEH sets. In addition, the categories related to abscisic acid signaling, which is known to work antagonistically to ethylene, GO:0009738 (abscisic acid-activated signaling pathway) and GO:0071215 (cellular response to abscisic acid stimulus), were also detected only in *A* with many inactivated genes by treatment. In contrast, the categories related to oxidative stress showed the strongest enrichment in *R* than others, suggesting higher intensity of oxidative stress in *C. rivularis* than other species.

### 3.7 VEHs related to meristem and their GO enrichment analysis

Among the GO categories enriched in four VEH sets, three categories were related to meristem activity: GO:0035266 (meristem growth), GO:0010075 (regulation of meristem growth), and GO:0048509 (meristem development) (**Table 1**). They were enriched only in VEH sets of *I_A_* and *I_R_*, but not in *A* and *R*, although *C. rivularis* can also produce ectopic meristems.

We analyzed the expression pattern of several known transcriptional factors which could be involved in ectopic meristem formation and development in *Cardamine* (**Fig. 3** and **Fig. S8**). Class I Knotted1-like homeobox (KNOX) transcription factors function to maintain shoot apical meristem activity in many different plant species (Vollbrecht et al., 1991; Long et al., 1996; Hay and Tsiantis, 2010). Importantly, the overexpression of *SHOOTMERISTEMLESS* (*STM*) and another KNOX gene, *Arabidopsis knotted 1*-like gene (*KNAT1*) are known to cause ectopic meristem formation on the leaf in *A*. *thaliana* (Chuck et al., 1996; Williams, 1998). Moreover, an *STM* ortholog is required for leaf vivipary in *Kalanchoë daigremontiana* (Garcês et al., 2007), a clonal propagation trait that is also observed in *C. rivularis* and *C. insueta*. As summarized in Fig. 3, orthologs of the four *A. thaliana* KNOXI genes, *STM*, *KNAT1, KNAT2 and KNAT6*, showed upregulated expression in all or any of *I_A_*, *I_R_* and *R*, but not in *A*. In addition, we also found that *PDF1* increased expression levels in *C. insueta* (both *I_A_* and *I_R_*) and *R*, which is exclusively detected in the L1 layer of shoot apical meristem throughout the shoot development of *Arabidopsis* (Abe et al., 1999). Three other transcription factor-encoding genes, *CUC2*, *CUC3* and *LAS*, which contribute to ectopic shoot apical meristem in tomato leaves (Rossmann et al., 2015), were induced in *R* and *I_R_* but not in *A*.

**Figure 3.**
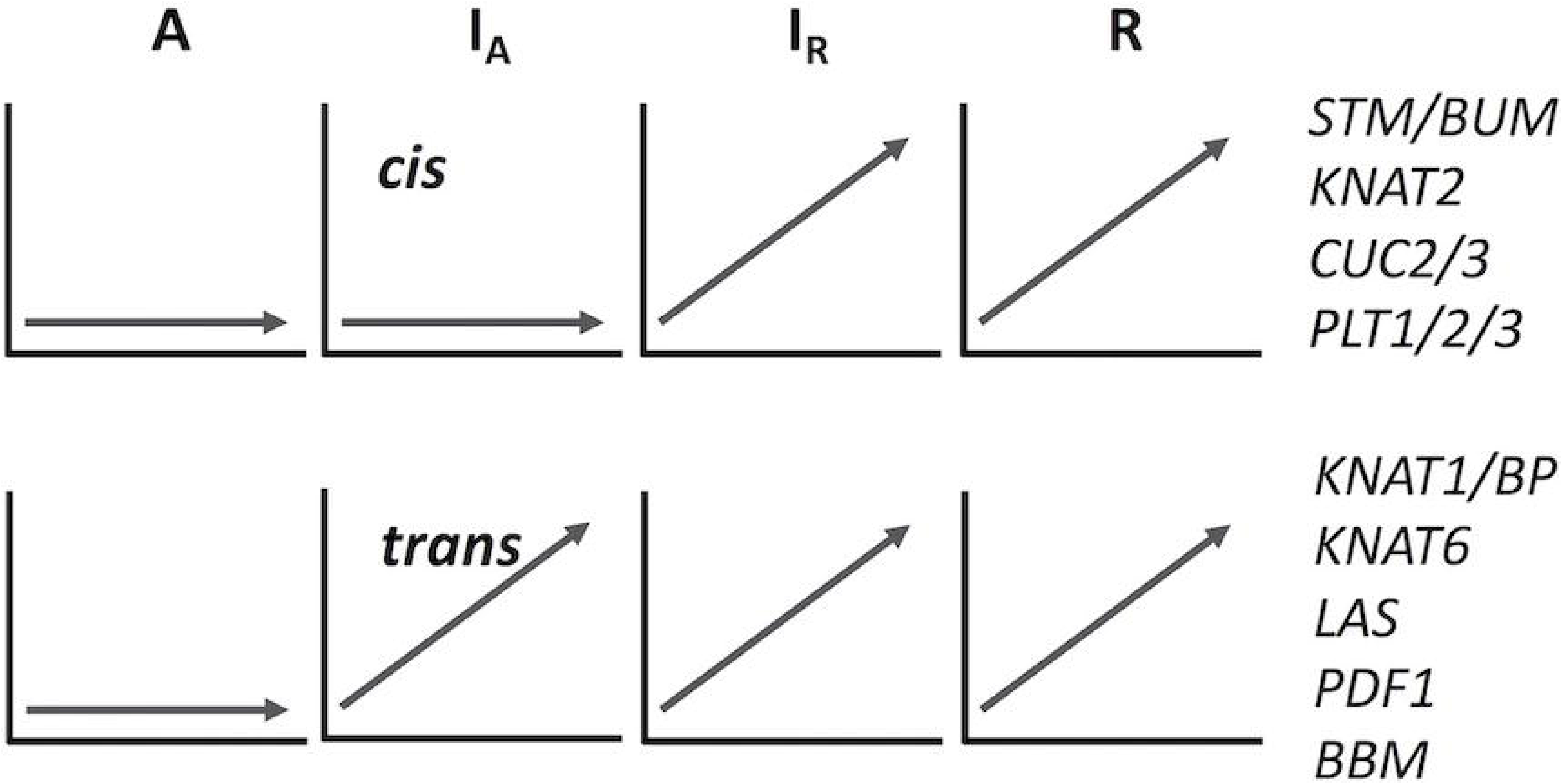
A schematic drawing of *cis*- and *trans*-regulation of key regulatory genes in meristem formation. The expression patterns of I_A_ homeologs are arbitrary categorized to *cis*- or *trans*-regulated according to the expression pattern of each homeolog. See Figure S8 for their original temporal expression patterns.

The expression of genes related to root apical meristem maintenance and formation showed similar patterns to those related to shoot apical meristem formation. Transcription factors with an AP2/ERF domain for the maintenance of root apical meristem (*PLT1*, *PLT2* and *PLT3*; (Drisch and Stahl, 2015)), were scarcely expressed in *A* but induced in others. For genes with similar function, *SHR* and *SCR*, expression level of *SHR* was increased in all four sets, while that of *SCR* was upregulated only temporarily and reverted in 24 hr. The expression of *WOX5*, a key factor to maintain the root stem cell (Sarkar et al., 2007), was very low in all sets, most probably due to the extremely limited expression area only at the quiescent center.

Many of the above-mentioned transcription factors contributing to meristem formation and maintenance are known to be related to or controlled by auxin, thus the transportation of auxin might be also involved in ectopic meristem development in *C. insueta* and *C. rivularis*. One of the auxin transporter genes, *PIN1* was induced by the treatment in all four sets soon after the start of submergence, but after 24 hr the high expression level was only retained in *I_A_*, *I_R_*, and *R*. On the other hand, the other auxin transporter genes *PIN3*, *PIN4*, and *PIN7* were temporarily induced by the treatment, but soon decreased among all sets. The peaks of expression of these three genes were at 4– 12 hr in A, 12–24 hr in R, and 8–24 hr in *I_A_* and *I_R_*, suggesting involvement in meristem formation and development. By the VEH enrichment analysis, GO:0060918 (auxin transport) and GO:0009926 (auxin polar transport) were enriched in *I_A_* and *I_R_*. In contrast, the other two auxin related GO categories, GO:0009850 (auxin metabolic process) and GO:0009851 (auxin biosynthetic process), were enriched in only two parents, *A* and *R*.

## 4 Discussion

### 4.1 Applicability of HomeoRoq to diverse ploidy levels

HomeoRoq was developed to classify genomic origins of RNA-Seq reads of allopolyploids consisting of two subgenomes (Akama et al., 2014), and has already been applied to *Arabidopsis kamchatica* (2*n* = 4*x* = 32; HHLL), an allotetraploid between two diploids of *Arabidopsis halleri* (2*n* = 2*x* = 16; HH) and *Arabidopsis lyrata* (2*n* = 2*x* = 16; LL). Here, we successfully applied HomeoRoq to another species with a different ploidy level. The average proportions of the reads mapped on the wrong genome in *C. amara* and *C. rivularis* samples were 1.1 ± 0.1% and 1.2 ± 0.1%, respectively (**Dataset S2**). This high accuracy is comparable to the evaluation of the *A. kamchatica* data, 1.23%– 1.64% (Akama et al., 2014; Kuo et al., 2020).

The proportion of *unclassified* reads in this study, which has the same matching rates on both parental genomes, was very close to that in the *A. kamchatica* study. In this study, 11.5 ± 2.0% of reads in *C. insueta* samples were *unclassified* on average, compared to 11.0% in *A. kamchatica* (Akama et al., 2014), suggesting a similar divergence level between subgenomes in the two cases. Considering the percentage of unclassified reads and the low misclassification rate with diploid progenitors, HomeoRoq can be applied to genomes of any ploidy level providing that the genome consists of two types of subgenome.

### 4.2 Total gene expression level of each subgenome is consistent with the chromosome number

The ratio of *A-origin* to *R-origin* reads in *C. insueta* was approximately 1:2. This result is consistent with the distribution of A-origin ratio showing a gentle peak at around 0.33 with a smooth decrease toward the edges (**Fig. 1**). This distribution indicates that expression ratios of most homeologs correlates with the copy number. A similar tendency could be found in other Brassicaceae allotetraploids (2, 5, 6). In the analysis of triploid banana (2*n* = 3*x* = 33; ABB), a hybrid between *Musa acuminata* (2*n* = 2*x* = 22; AA) and *M. balbisiana* (2*n* = 2*x* = 22; BB), the read proportion is distributed around 0.66 for the B alleles by 155 homeologs with rather high expression level detected by LC-MSMS as isoforms (van Wesemael et al., 2018). This could also be seen in hexaploid bread wheat consisting of three subgenomes, where 70% of genes showed balanced expression among homeologs (Ramírez-González et al., 2018). So far, this consistency between ploidy number and expression ratio looks like a general rule in many species with some exceptions like tetraploid cotton (Yoo et al., 2013).

In addition to the majority of genes that show balanced expression between homeologs, a limited proportion of genes show significant differential expression. Even though a direct comparison among studies is difficult due to different thresholding policies, the number of genes with unbalanced homeolog expression tends to be the minor fraction in many quantitative studies. Further studies should show whether a similar pattern is observed in even higher ploidy levels or other odd ploidies.

### 4.3 Limited number of homeolog pairs changed expression ratio in submergence condition

Though the number of homeologs with unbalanced expression is smaller than that with balanced expression, they could play a significant role in speciation of polyploid species, especially for achieving a combined trait from progenitors. A series of studies have reported that homeolog expression ratios can be changed depending on external environments (Bardil et al., 2011; Dong and Adams, 2011; Akama et al., 2014; Paape et al., 2016). Akama *et al*. evaluated the changes of the homeolog expression ratio of *A. kamchatica* after cold treatment (Akama et al., 2014). They reported that the homeolog expression ratios before and after cold treatment were highly correlated (*R*_2_ = 0.87), and only 1.11% of homeolog pairs statistically significantly changed in expression ratios in response to cold treatment (Akama et al., 2014). A similar result was reported for zinc treatment of *A. kamchatica*. The correlation of homeolog expression ratios between zinc treatment and control ranged from 0.89 to 0.94, and 0.3%–1.5% of homeologs significantly changed expression ratios after Zn treatment (Paape et al., 2016).

In this study using another Brassicaceae species, *C. insueta*, the correlation coefficients of A-origin ratios between 0 hr and the other time-points ranged from 0.68 to 0.82 (**Table S1**). The lowest correlation occurring between 2 hr and other time points may suggest that the initial reaction to the treatment had the strongest effect on gene expression. The overall high correlations among time points indicate that the expression ratios of most homeologs do not change considerably in response to treatment. Even though *C. rivularis* and *C. amara* show species-specific responses to submergence, leaf vivipary and submergence tolerance respectively, no specific expression preference or dominance of either progenitor was detected in the triploid. This suggests that transcriptional changes in only a limited number of homeologs, rather than genome-wide, might be responsible for the control of physiological change under submergence conditions.

### 4.4 Triploid inherited advantageous traits from progenitors

Only about 6% of the expressed genes were detected as VEH throughout the 96-hr treatment in each genome and subgenome, suggesting the criteria were fairly conservative. Among enriched GO categories are water stress related ones, particularly ethylene-response and fermentation. Fermentation metabolism in plants is important for submergence stress. We found more VEH genes in the fermentation-related categories in the diploid *C. amara* and the *amara*-derived subgenome of *C. insueta* than counterparts (Table 1). This suggests that *C. insueta* inherited the fermentation ability as a submergence response more largely from *C. amara* side. The ethylene signaling pathway should be stimulated in all three species as many related GO categories are found enriched in all VEH genes. However, the stress level seems to be variable according to the species as shown in the difference of enriched GO categories. In all of these ethylene related GO categories, *C. amara* had the strongest enrichment (i.e., highest number of VEH genes), and the enrichment in *amara*-derived subgenome was stronger than in *rivularis*-derived subgenome in *C. insueta*. These enrichment intensities should suggest that *C. amara* has higher acclimation ability to submergence through an activation of alcohol metabolic pathway and alteration in hormone signaling pathway and thus suffer from less oxidative stress as a result, as speculated by its habitat and a previous study (Shimizu-Inatsugi et al., 2016). In addition, in *C. insueta*, the contribution to the stress response of *I_A_* seems larger than that of *I_R_*, found as stronger enrichment in *I_A_* than in *I_R_*.

GO enrichment analysis with VEH genes also showed three GO categories related to meristem, GO:0035266 (meristem growth), GO:0010075 (regulation of meristem growth) and GO:0048509 (meristem development). They were only enriched in the VEH sets of *I_A_* and *I_R_*, but not above the significance threshold in two parents, despite the fact that *C. rivularis* also produces plantlets on the leaf by the activation of ectopic meristems. This might imply that the ability to form ectopic plantlets in response to submergence is enhanced in the triploid *C. insueta* compared to the diploid *C. rivularis*. Considering the disadvantage in sexual reproduction due to the odd ploidy, effective vegetative propagation through plantlets might have been critically important for *C. insueta*.

The expression pattern of known key regulatory genes that function to maintain meristem activity showed two typical patterns, as shown in **Fig. 3** and **Fig S8**. Expression of these genes was upregulated in *C. rivularis* (*R*, **Fig. 3**) but not in *C. amara* (*A*, **Fig. 3**) in response to submergence. Expression of these genes was also upregulated in the *C. insueta* subgenome *I_R_*, but followed two different patterns in the *I_A_* subgenome. These patterns could be categorized as either non-induced, similar to *C. amara*, or induced, similar to *C. rivularis*, suggesting that non-induced homeologs could be *cis*-regulated by *I_A_*, and induced homeologs could be *trans*-regulated by *I_R_*. One possibility is that this difference reflects the developmental timing of gene expression during meristem formation. For example, the *cis*-regulated genes *STM* and *CUC2* are expressed earlier during embryogenesis in *A. thaliana* than the *trans*-regulated genes *KNAT6* and *KNAT1/BP* (Hay and Tsiantis, 2010). This variation might imply a regulatory relationship among these genes in the gene regulatory network controlling plantlet formation in *C. insueta* leaves. This type of information might provide insights that warrant further study into the molecular mechanism of leaf vivipary in *C. rivularis* and *C. insueta*.

## 5 Acknowledgments

We would like to thank Walter Brücker for the support in the fieldwork in Urnerboden, Martin Lysak, Terezie Mandáková, Karol Marhold, Judita Zozomová-Lihová, Pamela Soltis and Douglas Soltis for discussion. Data analysis was partially performed on National Institute of Genetics (NIG) at Research Organization of Information and Systems (ROIS), Japan.

## 6 Author Contributions

JSun analysed data. JSun and RSI wrote the manuscript. RSI, AH, KKS and JSese refined the manuscript. HH performed experiment. AH, KKS, JSese supervised the project. All authors read, corrected, and approved the manuscript.

## 7 Conflict of Interest

The authors declare that the research was conducted in the absence of any commercial or financial relationships that could be construed as a potential conflict of interest.

## 8 Contribution to the Field Statement

In the research of genome evolution, a newborn species offers a unique and interesting case to study how the genome evolves during speciation and adaptation. A *Brassicaceae* plant, *Cardamine insueta*, which was born in a small village in Swiss Alps in the 20_th_ century, applies to this. *C. insueta* is generated by the hybridization and genome duplication between two closely related progenitor species, which owned their respective habitats. This plant has adapted to a new habitat covering the intermediate area between the progenitor species. How could it achieve this in the course of speciation? The usage of the genes inherited from two progenitors should have the clue to answer this question. We studied the gene expression pattern of *C. insueta* to analyze how it regulates the two types of the same gene inherited from its diploid progenitors, and found that it conserves the gene expression pattern of the advantageous progenitor according to different traits. For the trait of submergence tolerance, it exploits the pattern of one progenitor having the trait. For another trait, clonal propagation, it exploits the pattern of the other progenitor having this trait. This result contributes to our understanding of speciation, and how various genes are affected in a speciation process.

## 9 Funding

This study was funded by the Swiss National Science Foundation to RSI (Marie-Heim Högtlin grant) and KKS (31003A_182318); by University Research Priority Programs, Evolution in Action of the University of Zurich to RSI and KKS; by the Human Frontier Science Program to KKS, AH, and JSese; by the Japan Science and Technology Agency, Core Research for Evolutionary Science and Technology grant number JPMJCR16O3, Japan; and by KAKENHI grant numbers 16H06469 to JSese, KKS, and JSun.

## 12 Supplementary Material

Supplementary_Material.pdf (Table S1, Fig. S1 to Fig. S8)

**SI dataset 1** Videos for visualizing of plantlet initiation of *C. insueta* Video file (MP4) visualizes the plantlet initiation of *C. insueta* at the incubator under the standard condition with the 16 hr light and 8 hr dark. Leaflet were detached form *C. insueta* individuals and floated on the water in a beaker. Photos were taken between May 16, 2018 and June 7, 2018, with 90 minutes intervals. Photos taken at daylight were concatenated into video.

**SI dataset 2** Statistics of RNA-Seq data processing An Excel format file that contains the statistics of RNA-Seq data processing with HomeoRoq pipeline. Sheet 1: Number of read pairs before and after quality controls with Trimmomatic; Sheet 2: Number of reads that were mapped onto A-genome and R-genome, and number of reads that were classified into A-origin, R-origin, and unclassified reads with HomeoRoq.

**SI dataset 3** FPKM and CV of variably expressed homeologs An Excel format file that contains gene names, averages of log10-transformed FPKM, and coefficient of variation (CV) of log10-transformed FPKM of variably expressed homeologs (VEHs). Sheet 1: VEHs of IA samples; Sheet 2: VEHs of IR samples; Sheet 3: VEHs of C. amara samples; Sheet 4: VEHs of C. rivularis samples.

**SI dataset 4** Enriched GO terms of variably expressed homeologs An Excel format file that contains gene ontology (GO) enrichment analysis results of VEHs. Sheet 1: Summarization of GO enrichment analysis results. Values in cells represent negative log10(q-value); Sheet IA: GO enrichment analysis result of VEHs of I_A_ samples; Sheet IR: GO enrichment analysis result of VEHs of I_R_ samples; Sheet A: GO enrichment analysis result of VEHs of *C. amara* samples; Sheet R: GO enrichment analysis result of VEHs of *C. rivularis* samples.

**SI dataset 5** Relative gene expression levels of the genes of the category GO:0009414 (response to water deprivation).

## 13 Data Availability Statement

The datasets generated for this study can be found in DNA Data Bank of Japan (DDBJ) Sequence Read Archive (DRA), www.ddbj.nig.ac.jp [accession no. PRJDB9426].

## Notes

### Competing Interest Statement

The authors have declared no competing interest.

## References

Abbott, R., Albach, D., Ansell, S., Arntzen, J. W., Baird, S. J. E., Bierne, N., et al. (2013). Hybridization and speciation. J. Evol. Biol. 26, 229–246. doi:10.1111/j.1420-9101.2012.02599.x

Abbott, R., and Andrew, J. (2004). Origins, establishment and evolution of new polyploid species: Senecio cambrensis and S. eboracensis in the British Isles. Biol. J. Linn. Soc. 82, 467–474. doi:10.1111/j.1095-8312.2004.00333.x

Abe, M., Takahashi, T., and Komeda, Y. (1999). Cloning and characterization of an L1 layer-specific gene in *Arabidopsis thaliana*. Plant Cell Physiol. 40, 571–580. doi:10.1093/oxfordjournals.pcp.a029579

Ainouche, M. L., Baumel, A., and Salmon, A. (2004). Spartina anglica C. E. Hubbard: a natural model system for analysing early evolutionary changes that affect allopolyploid genomes. Biol. J. Linn. Soc. 82, 475–484. doi:10.1111/j.1095-8312.2004.00334.x

Akama, S., Shimizu-Inatsugi, R., Shimizu, K. K., and Sese, J. (2014). Genome-wide quantification of homeolog expression ratio revealed nonstochastic gene regulation in synthetic allopolyploid Arabidopsis. Nucleic Acids Res. 42, 1–15. doi:10.1093/nar/gkt1376

Akiyama, R., Sun, J., Hatakeyama, M., Lischer, H. E. L., Briskine, R. V., Hay, A., et al. (2019). Fine-scale ecological and transcriptomic data reveal niche differentiation of an allopolyploid from diploid parents in *Cardamine*. bioRxiv, 600783. doi:10.1101/600783

Alabadí, D., Oyama, T., Yanovsky, M. J., Harmon, F. G., Más, P., and Kay, S. A. (2001). Reciprocal regulation between TOC1 and LHY/CCA1 within the *Arabidopsis* circadian clock. Science 293, 880–883. doi:10.1126/science.1061320

Bardil, A., de Almeida, J. D., Combes, M. C., Lashermes, P., and Bertrand, B. (2011). Genomic expression dominance in the natural allopolyploid *Coffea arabica* is massively affected by growth temperature. New Phytol. 192, 760–774. doi:10.1111/j.1469-8137.2011.03833.x

Bolger, A. M., Lohse, M., and Usadel, B. (2014). Trimmomatic: A flexible trimmer for Illumina sequence data. Bioinformatics 30, 2114–2120. doi:10.1093/bioinformatics/btu170

Bretagnolle, F., and Thompson J. D. (1995). Gametes with the somatic chromosome number: mechanisms of their formation and role in the evolution of autopolyploid plants. New Phytol. 129, 1–22. doi:10.1111/j.1469-8137.1995.tb03005.x

Chao, Q., Rothenberg, M., Solano, R., Roman, G., Terzaghi, W., and Ecker†, J. R. (1997). Activation of the ethylene gas response pathway in Arabidopsis by the nuclear protein ETHYLENE-INSENSITIVE3 and related proteins. Cell 89, 1133–1144. doi:10.1016/S0092-8674(00)80300-1

Chuck, G., Lincoln, C., and Hake, S. (1996). *KNAT1* induces lobed leaves with ectopic meristems when overexpressed in Arabidopsis. Plant Cell 8, 1277–1289. doi:10.1105/tpc.8.8.1277

Comai, L. (2005). The advantages and disadvantages of being polyploid. Nat. Rev. Genet. 6, 836–846. doi:10.1038/nrg1711

Czechowski, T., Stitt, M., Altmann, T., and Udvardi, M. K. (2005). Genome-wide identification and testing of superior reference genes for transcript normalization. Society 139, 5–17. doi:10.1104/pp.105.063743.1

Dickinson, T. A. (1978). Epiphylly in angiosperms. Bot. Rev. 44, 181–232. doi:10.1007/BF02919079

Dobin, A., Davis, C. A., Schlesinger, F., Drenkow, J., Zaleski, C., Jha, S., et al. (2013). STAR: ultrafast universal RNA-seq aligner. Bioinformatics 29, 15–21. doi:10.1093/bioinformatics/bts635

Dong, S., and Adams, K. L. (2011). Differential contributions to the transcriptome of duplicated genes in response to abiotic stresses in natural and synthetic polyploids. New Phytol. 190, 1045–1057. doi:10.1111/j.1469-8137.2011.03650.x

Drisch, R. C., and Stahl, Y. (2015). Function and regulation of transcription factors involved in root apical meristem and stem cell maintenance. Front. Plant Sci. 6, 1–8. doi:10.3389/fpls.2015.00505

Gan, X., Hay, A., Kwantes, M., Haberer, G., Hallab, A., Ioio, R. Dello, et al. (2016). The *Cardamine hirsuta* genome offers insight into the evolution of morphological diversity. Nat. plants 2, 16167. doi:10.1038/nplants.2016.167

Garcês, H. M. P., Champagne, C. E. M., Townsley, B. T., Park, S., Malhó, R., Pedroso, M. C., et al. (2007). Evolution of asexual reproduction in leaves of the genus *Kalanchoë*. Proc. Natl. Acad. Sci. U. S. A. 104, 15578–15583. doi:10.1073/pnas.0704105104

Hay, A., and Tsiantis, M. (2010). KNOX genes: versatile regulators of plant development and diversity. Development 137, 3153–3165. doi:10.1242/dev.030049

Howard, H. W. (1948). Chromosome number of *Cardamine pratensis*. Nature 161, 277–277. doi:10.1038/161277a0

Husband, B. C. (2004). The role of triploid hybrids in the evolutionary dynamics of mixed-ploidy populations. Biol. J. Linn. Soc. 82, 537–546. doi:10.1111/j.1095-8312.2004.00339.x

Hussein, F. (1948). Chromosome number of *Cardamine pratensis*. Nature 161, 1015–1015. doi:10.1038/1611015a0

Jerry A, C., and H. Allen, O. (2004). Speciation. Sunderland, Mass: Sinauer Associates

Kuo, T. C. Y., Hatakeyama, M., Tameshige, T., Shimizu, K. K., and Sese, J. (2020). Homeolog expression quantification methods for allopolyploids. Brief. Bioinform. 21, 395–407. doi:10.1093/bib/bby121

Li, H., Handsaker, B., Wysoker, A., Fennell, T., Ruan, J., Homer, N., et al. (2009). The Sequence Alignment/Map format and SAMtools. Bioinformatics 25, 2078–2079. doi:10.1093/bioinformatics/btp352

Long, J. A., Moan, E. I., Medford, J. I., and Barton, M. K. (1996). A member of the KNOTTED class of homeodomain proteins encoded by the *STM* gene of *Arabidopsis*. Nature 379, 66–69. doi:10.1038/379066a0

Mable, B. K. (2003). Breaking down taxonomic barriers in polyploidy research. Trends Plant Sci. 8, 582–590. doi:10.1016/j.tplants.2003.10.006

Mandáková, T., Kovařík, A., Zozomová-Lihová, J., Shimizu-Inatsugi, R., Shimizu, K. K., Mummenhoff, K., et al. (2013). The More the Merrier: Recent Hybridization and Polyploidy in *Cardamine*. Plant Cell 25, 3280–3295. doi:10.1105/tpc.113.114405

Mandáková, T., Marhold, K., and Lysak, M. A. (2014). The widespread crucifer species *Cardamine flexuosa* is an allotetraploid with a conserved subgenomic structure. New Phytol. 201, 982–992. doi:10.1111/nph.12567

Mason, A. S., and Pires, J. C. (2015). Unreduced gametes: Meiotic mishap or evolutionary mechanism? Trends Genet. 31, 5–10. doi:10.1016/j.tig.2014.09.011

Paape, T., Hatakeyama, M., Shimizu-Inatsugi, R., Cereghetti, T., Onda, Y., Kenta, T., et al. (2016). Conserved but attenuated parental gene expression in allopolyploids: constitutive zinc hyperaccumulation in the allotetraploid *Arabidopsis kamchatica*. Mol. Biol. Evol. 33, 2781–2800. doi:10.1093/molbev/msw141

Planet, E., Attolini, C. S. O., Reina, O., Flores, O., and Rossell, D. (2012). htSeqTools: High-throughput sequencing quality control, processing and visualization in R. Bioinformatics 28, 589–590. doi:10.1093/bioinformatics/btr700

Ramírez-González, R. H., Borrill, P., Lang, D., Harrington, S. A., Brinton, J., Venturini, L., et al. (2018). The transcriptional landscape of polyploid wheat. Science 361, eaar6089. doi:10.1126/science.aar6089

Ramsey, J., and Schemske, D. W. (1998). Pathways, mechanisms, and rates of polyploid formation in flowering plants. Annu. Rev. Ecol. Syst. 29, 467–501. doi:10.1146/annurev.ecolsys.29.1.467

Rossmann, S., Kohlen, W., Hasson, A., and Theres, K. (2015). Lateral suppressor and Goblet act in hierarchical order to regulate ectopic meristem formation at the base of tomato leaflets. Plant J. 81, 837–848. doi:10.1111/tpj.12782

Salisbury, E. J. (1965). The reproduction of *Cardamine pratensis* L. and *Cardamine palustris* Peterman particularly in relation to their specialized foliar vivipary, and its deflexion of the constraints of natural selection. Proc. R. Soc. London. Ser. B. Biol. Sci. 163, 321–342. doi:10.1098/rspb.1965.0072

Sarkar, A. K., Luijten, M., Miyashima, S., Lenhard, M., Hashimoto, T., Nakajima, K., et al. (2007). Conserved factors regulate signalling in *Arabidopsis thaliana* shoot and root stem cell organizers. Nature 446, 811–814. doi:10.1038/nature05703

Shimizu-Inatsugi, R., Terada, A., Hirose, K., Kudoh, H., Sese, J., and Shimizu, K. K. (2016). Plant adaptive radiation mediated by polyploid plasticity in transcriptomes. Mol. Ecol. 26, 193–207. doi:10.1111/mec.13738

Shin, H., Shannon, C. P., Fishbane, N., Ruan, J., Zhou, M., Balshaw, R., et al. (2014). Variation in RNA-Seq transcriptome profiles of peripheral whole blood from healthy individuals with and without globin depletion. PLoS One 9, 1–11. doi:10.1371/journal.pone.0091041

Smith, J. E. (1825). English Flora. III. Flora.

Solano, R., Stepanova, A., Chao, Q., and Ecker, J. R. (1998). Nuclear events in ethylene signaling: a transcriptional cascade mediated by ETHYLENE-INSENSITIVE3 and ETHYLENE-RESPONSE-FACTOR1. Genes Dev. 12, 3703–14. doi:10.1101/gad.12.23.3703

Soltis, D. E., Soltis, P. S., Pires, J. C., Kovarik, A., Tate, J. A., and Mavrodiev, E. (2004). Recent and recurrent polyploidy in *Tragopogon* (Asteraceae): cytogenetic, genomic and genetic comparisons. Biol. J. Linn. Soc. 82, 485–501. doi:10.1111/j.1095-8312.2004.00335.x

Soltis, P. S., and Soltis, D. E. (2009). The role of hybridization in plant speciation. Annu. Rev. Plant Biol. 60, 561–588. doi:10.1146/annurev.arplant.043008.092039

Tayalé, A., and Parisod, C. (2013). Natural pathways to polyploidy in plants and consequences for genome reorganization. Cytogenet. Genome Res. 140, 79–96. doi:10.1159/000351318

Urbanska-Worytkiewicz, K., and Landolt, E. (1974a). Biosystematic investigations in *Cardamine pratensis* L.s.I. I. Diploid taxa from Central Europe and their fertility relationships. Berichte des Geobot. Instituts der ETH Stift. Rübel 42, 42–139

Urbanska-Worytkiewicz, K., and Landolt, E. (1974b). Hybridation naturelle entre *Cardamine rivularis* SCHUR et *C. amara* L., ses aspects cytologiques et écologiques. Act. Soc. Helv. Sci. Nat., 89–90

Urbanska, K. M. (1977). Reproduction in natural triploid hybrids (2n=24) between *Cardamine rivularis* Schur and *C. amara* L. *er. Geobot*. Inst. ETH Stift. Rubel 44, 42–85

Urbanska, K. M., Hurka, H., Landolt, E., Neuffer, B., and Mummenhoff, K. (1997). Hybridization and evolution in *Cardamine* (*Brassicaceae*) at Urnerboden, Central Switzerland: Biosystematic and molecular evidence. Plant Syst. Evol. 204, 233–256. doi:10.1007/BF00989208

Urbaska-Worytkiewicz, K., and Landolt, E. (1972). Natürliche Bastarde zwischen *Cardamine amara* L. und *C. rivularis* Schur aus den Schweizer Alpen. Ber. Geobot. Inst. ETH. Stift. Rübel 41, 88–101. doi:10.5169/seals-377675

van Wesemael, J., Hueber, Y., Kissel, E., Campos, N., Swennen, R., and Carpentier, S. (2018). Homeolog expression analysis in an allotriploid non-model crop via integration of transcriptomics and proteomics. Sci. Rep. 8, 1353. doi:10.1038/s41598-018-19684-5

Vollbrecht, E., Veit, B., Sinha, N., and Hake, S. (1991). The developmental gene *Knotted-1* is a member of a maize homeobox gene family. Nature 350, 241–243. doi:10.1038/350241a0

Williams, R. W. (1998). Plant homeobox genes: many functions stem from a common motif. BioEssays 20, 280–282. doi:10.1002/(SICI)1521-1878(199804)20:4<280::AID-BIES2>3.0.CO;2-U

Yamaguchi, R., Imoto, S., Yamauchi, M., Nagasaki, M., Yoshida, R., Shimamura, T., et al. (2008). Predicting differences in gene regulatory systems by state space models. Genome Inform. 21, 101–113. doi:10.11234/gi1990.21.101

Yoo, M.-J., Szadkowski, E., and Wendel, J. F. (2013). Homoeolog expression bias and expression level dominance in allopolyploid cotton. Heredity (Edinb). 110, 171–180. doi:10.1038/hdy.2012.94

Yu, G., Wang, L.-G., Han, Y., and He, Q.-Y. (2012). clusterProfiler: an R Package for Comparing Biological Themes Among Gene Clusters. Omi. A J. Integr. Biol. 16, 284–287. doi:10.1089/omi.2011.0118

Zhao, J., Yang, F., Feng, J., Wang, Y., Lachenbruch, B., Wang, J., et al. (2017). Genome-wide constitutively expressed gene analysis and new reference gene selection based on transcriptome data: a case study from poplar/canker disease interaction. Front. Plant Sci. 8, 1–13. doi:10.3389/fpls.2017.01876

Zozomová-Lihová, J., Krak, K., Mandáková, T., Shimizu, K. K., Španiel, S., Vít, P., et al. (2014). Multiple hybridization events in *Cardamine* (Brassicaceae) during the last 150 years: revisiting a textbook example of neoallopolyploidy. Ann. Bot. 113, 817–830. doi:10.1093/aob/mcu012

