## Supplemental Table and Supplemental Figures for "A recently formed triploid *Cardamine insueta* inherits leaf vivipary and submergence tolerance traits of parents"

### *Supplementary Material*

#### Supplementary Table S1

**Pearson correlation coefficients of A-origin ratios.** Pearson correlation coefficients of A-origin ratios between two *C. insueta* samples among the nine time points.

| | 2 hr | 4 hr | 8 hr | 12 hr | 24 hr | 48 hr | 72 hr | 96 hr |
|---|---|---|---|---|---|---|---|---|
| 0 hr | 0.663 | 0.807 | 0.689 | 0.707 | 0.757 | 0.791 | 0.794 | 0.732 |
| 2 hr | | 0.685 | 0.672 | 0.693 | 0.716 | 0.722 | 0.697 | 0.639 |
| 4 hr | | | 0.745 | 0.762 | 0.778 | 0.800 | 0.811 | 0.747 |
| 8 hr | | | | 0.833 | 0.759 | 0.773 | 0.749 | 0.695 |
| 12 hr | | | | | 0.787 | 0.798 | 0.771 | 0.719 |
| 24 hr | | | | | | 0.839 | 0.820 | 0.735 |
| 48 hr | | | | | | | 0.839 | 0.760 |
| 72 hr | | | | | | | | 0.805 |

**Supplementary Figure S1**

**Habitats and relations among the three *Cardamine* species.** (***A***) *C. insueta* (triploid) was naturally formed by hybridization between *C. amara* (diploid) and *C. rivularis* (diploid) 100-150 years ago in their natural habitat in Swiss Alps. (***B***) Conceptual indication of the habitats of three species at Urnerboden: *C. amara* prefers wet habitats along waterside; *C. rivularis* prefers meadow; and allotriploid *C. insueta* can be found between them.

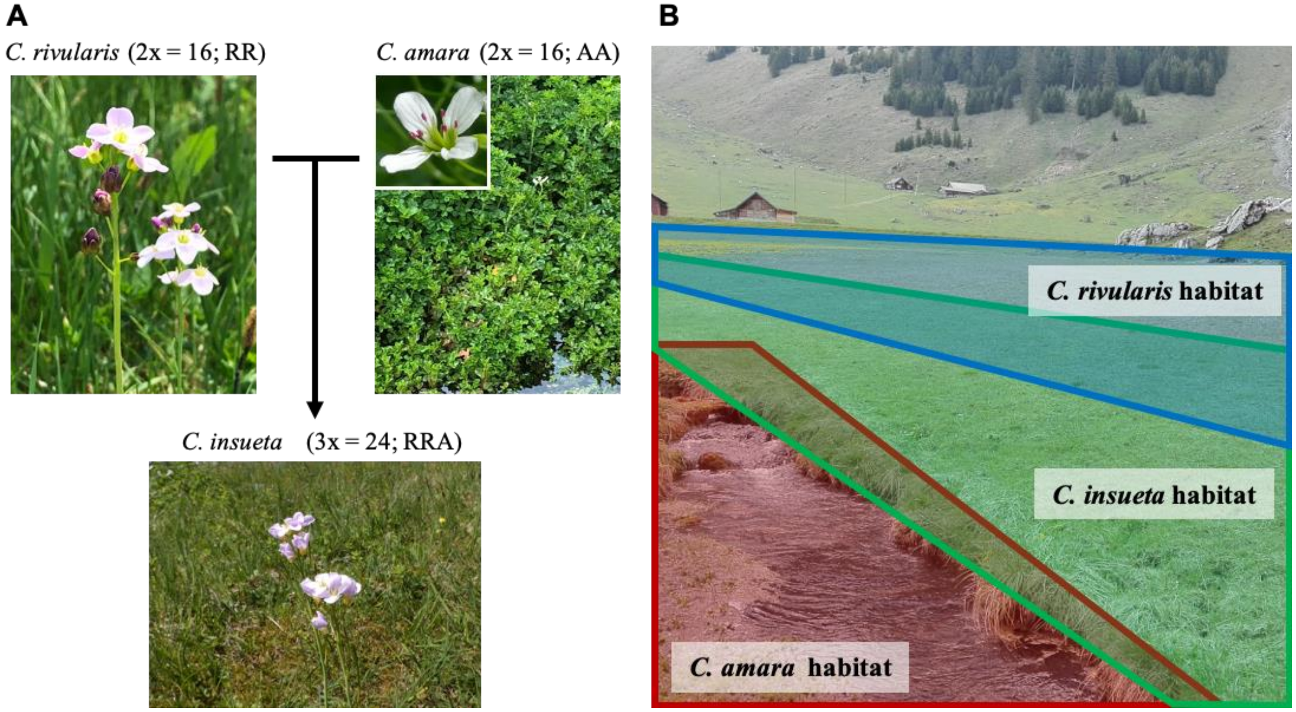

**Supplementary Figure S2**

**Leaf vivipary in a representative leaf of *C. rivularis*.** A new plantlet was first visible 96 hours after submergence (circled, shown in close-up below). Plantlets initiated from dormant shoot meristems (shoot meristem with visible leaf shown). Shoot growth was detected 8 hours after submergence, followed by root initiation (16 hours). The shoot and root poles appeared to fuse (72 hours), followed by shoot and root growth to produce a plantlet (96 hours).

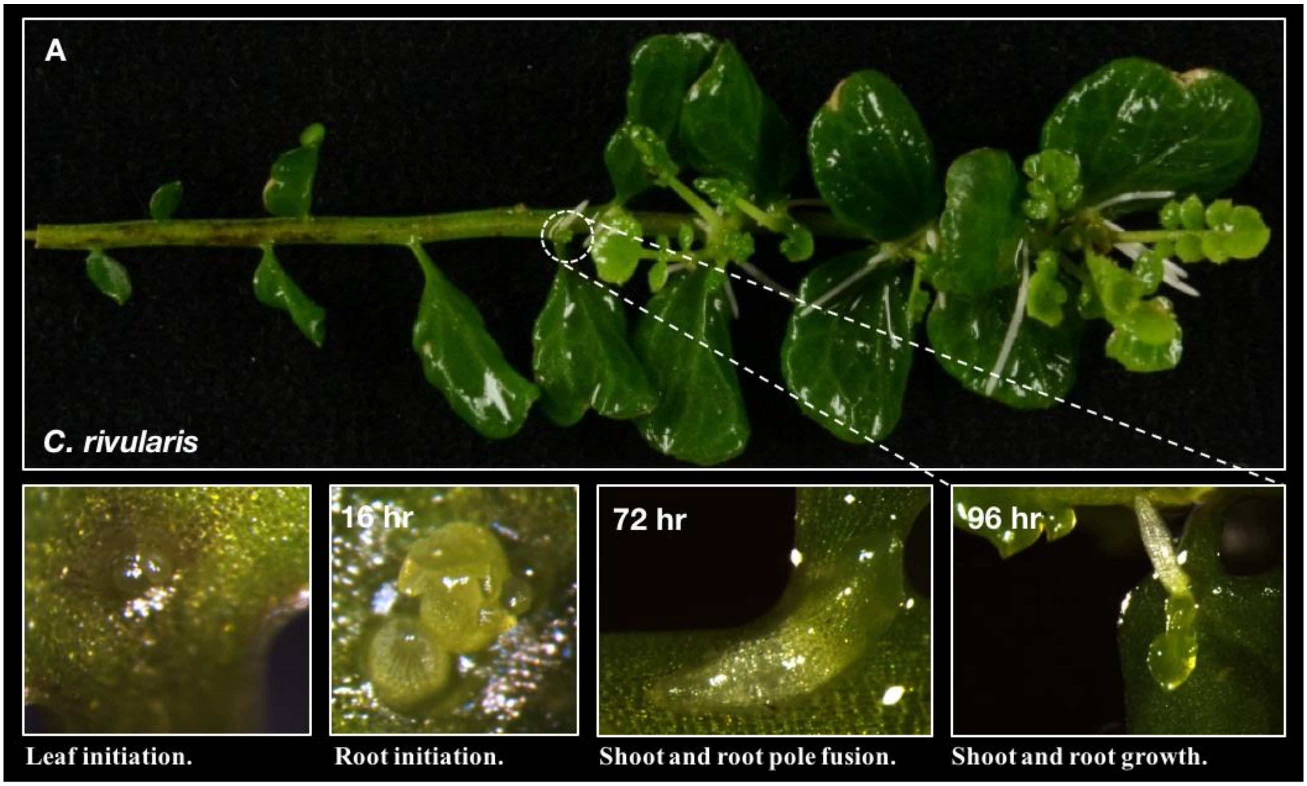

**Supplementary Figure S3**

**Overview of homeolog expression analysis.** (***A***) To monitor homeolog expression profiles during submergence responses, we sequenced 27 RNA samples extracted from the leaf of the three species (*C. insueta*, *C. amara*, and *C. rivularis*) at the nine time points after the start of submergence. (***B***) A-genome and R-genome were assembled from RNA-Seq reads of the nine *C. amara* and *C. rivularis* samples, respectively. (***C***) Homeolog-specific expression was quantified using HomeoRoq pipeline. For each *C. insueta* sample, reads were mapped onto both A-genome and R-genome. Then, homeolog-specific reads were classified into *A-origin*, *R-origin* and unclassified (with the same mismatch rates on A and R) according to the number of mismatches on the two mapping results. After classification, the read count data, FPKM (fragments per kilobase of exon per million reads mapped), and A-origin ratio were calculated from the results.

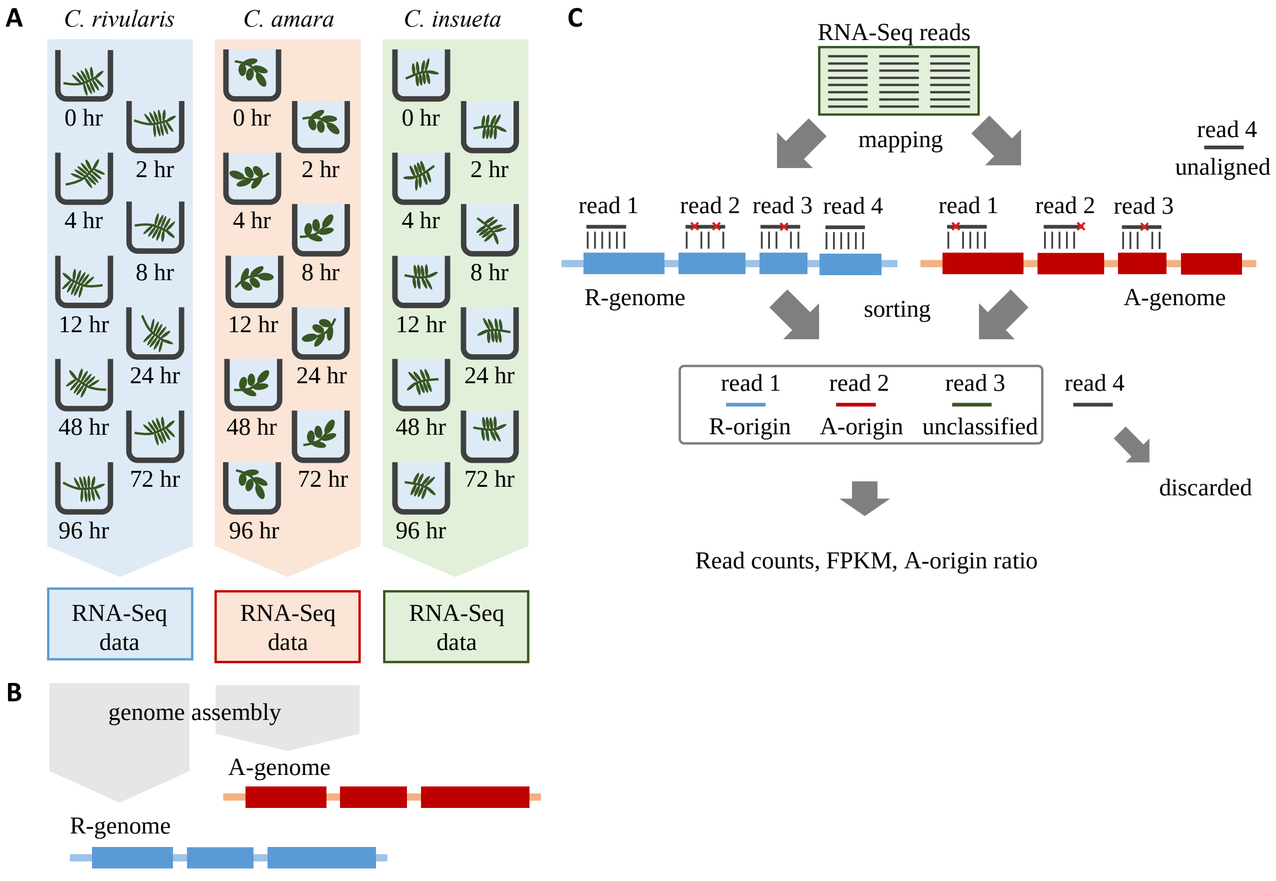

**Supplementary Figure S4**

**Expression ratio between A- and R-homeologs during submergence treatment in the triploid *C. insueta*.** Each dot shows the relation between the $\log_{10}$-transformed A-origin and R-origin reads of a homeolog pair at nine time points. Only the homeolog pairs with FPKM > 1.0 in either $I_A$ or $I_R$ samples are shown. The orange line represents the ratio A:R=1:2.

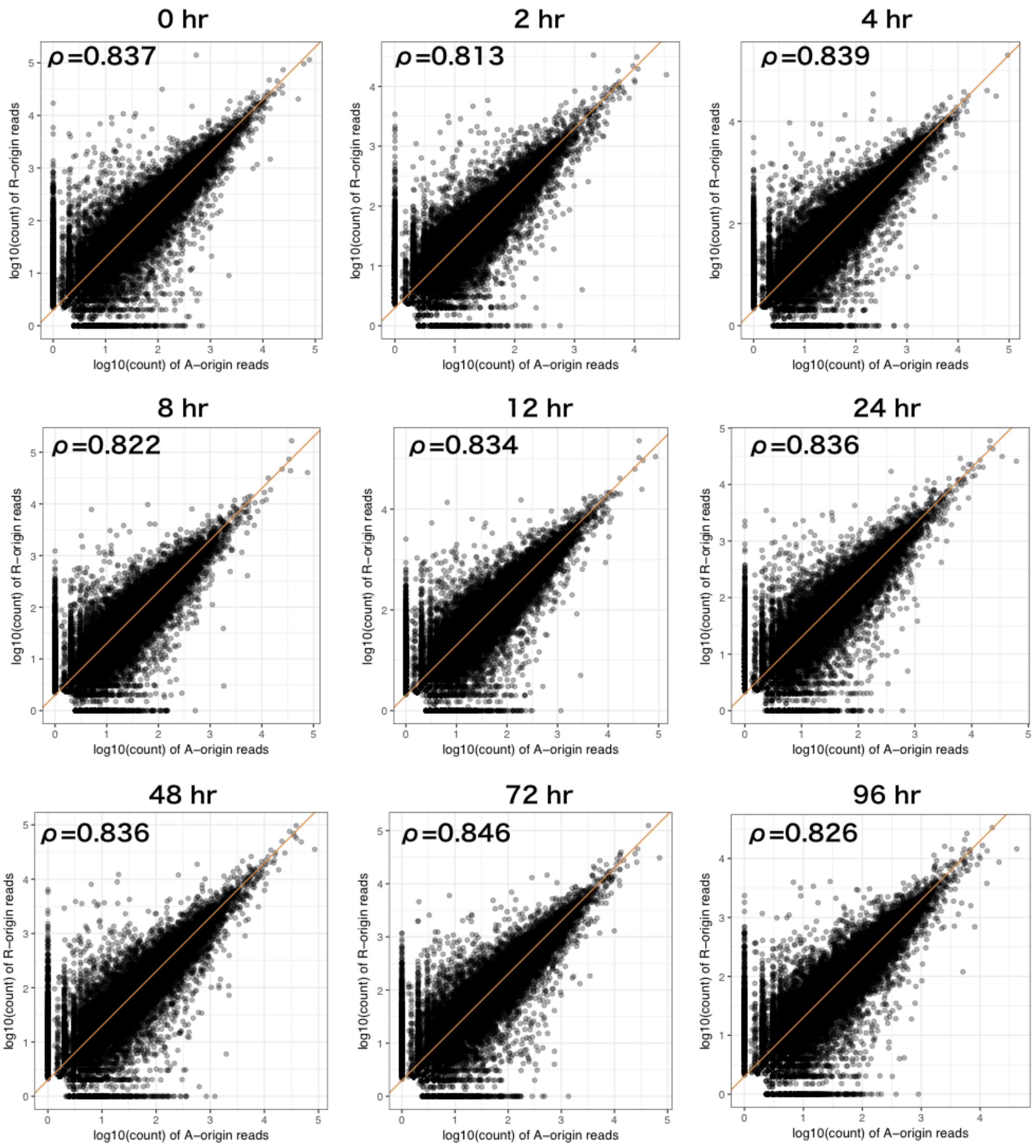

**Supplementary Figure S5**

**Distributions of A-origin ratio of *C. insueta* at nine time points.** The width of the bin is 0.025 in these histograms. The vertical orange lines indicate one-third of A-origin ratios. The number of homeologs for plotting histograms is shown in the title of each histogram.

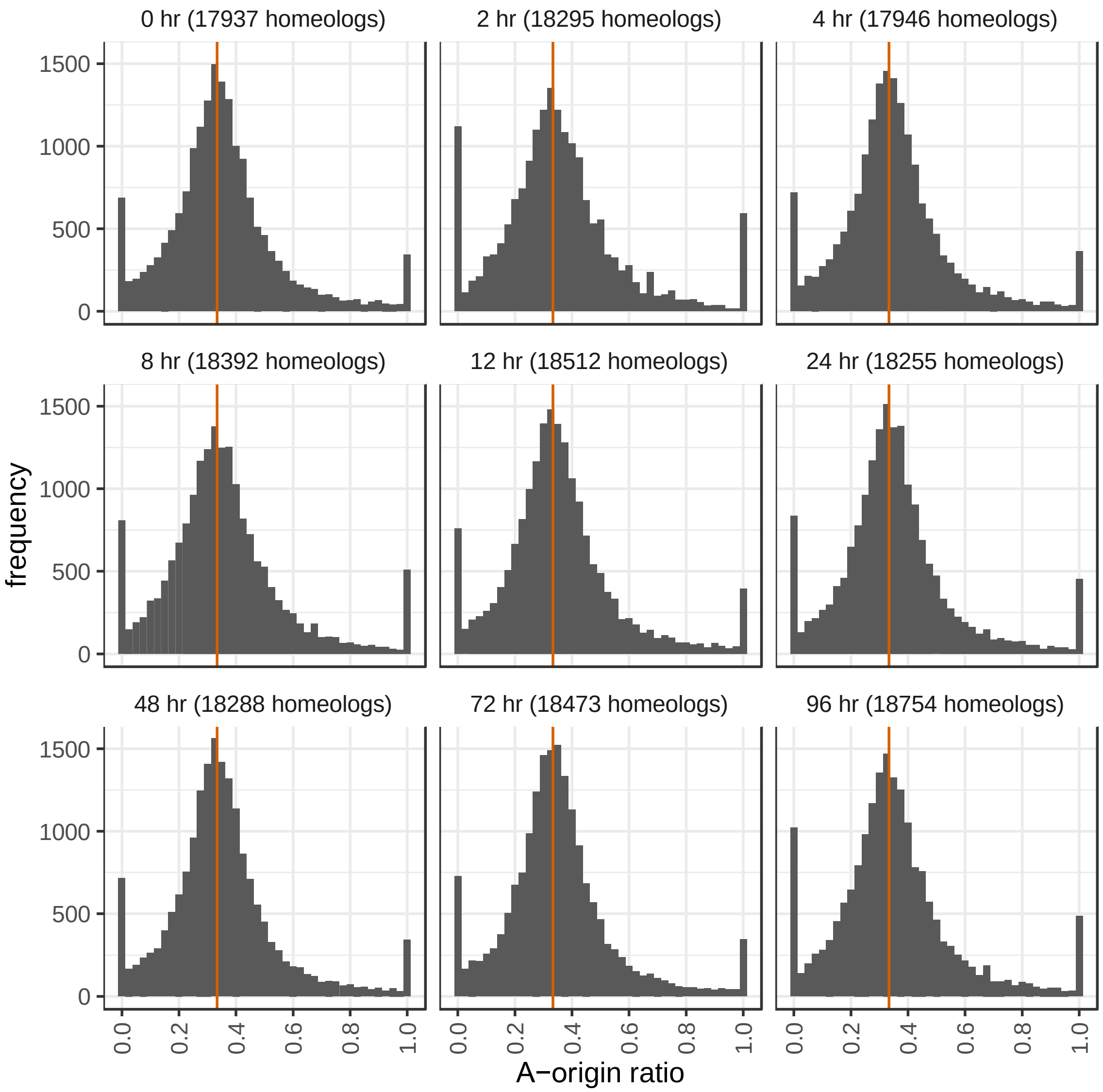

**Supplementary Figure S6**

**Time-course changes of homeolog expression.** Expression profiles of *ERF1* and *CCA1*. Top panels represent expression of the four homeologs in the IA, IR, *C. amara*, and *C. rivularis* samples.

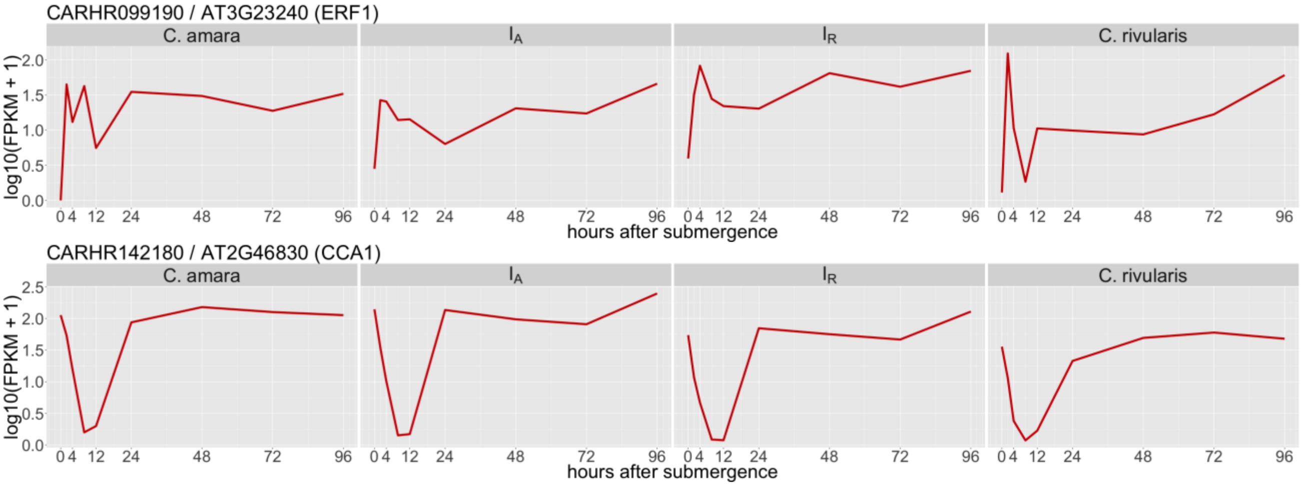

**Supplementary Figure S7**

**Overlaps of the number of VEH genes/homeologs of four genomes/subgenomes.**

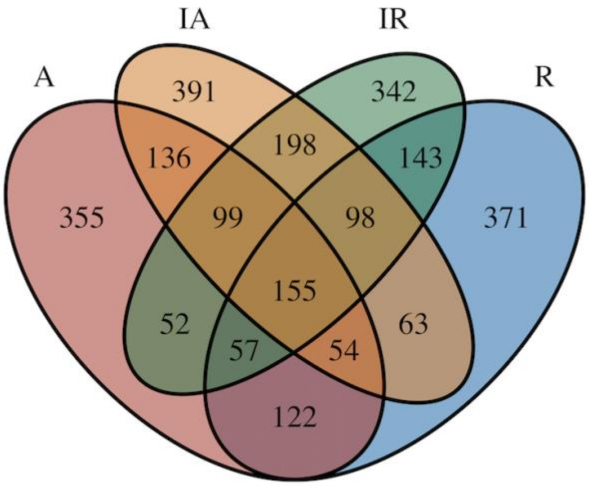

#### Supplementary Figure S8

**Time-course changes of homeolog expression of plantlet associated homeologs.** Expression profiles of known meristem associated homeologs are shown in line charts. Red line indicates that the gene/homeolog was identified as VEH.

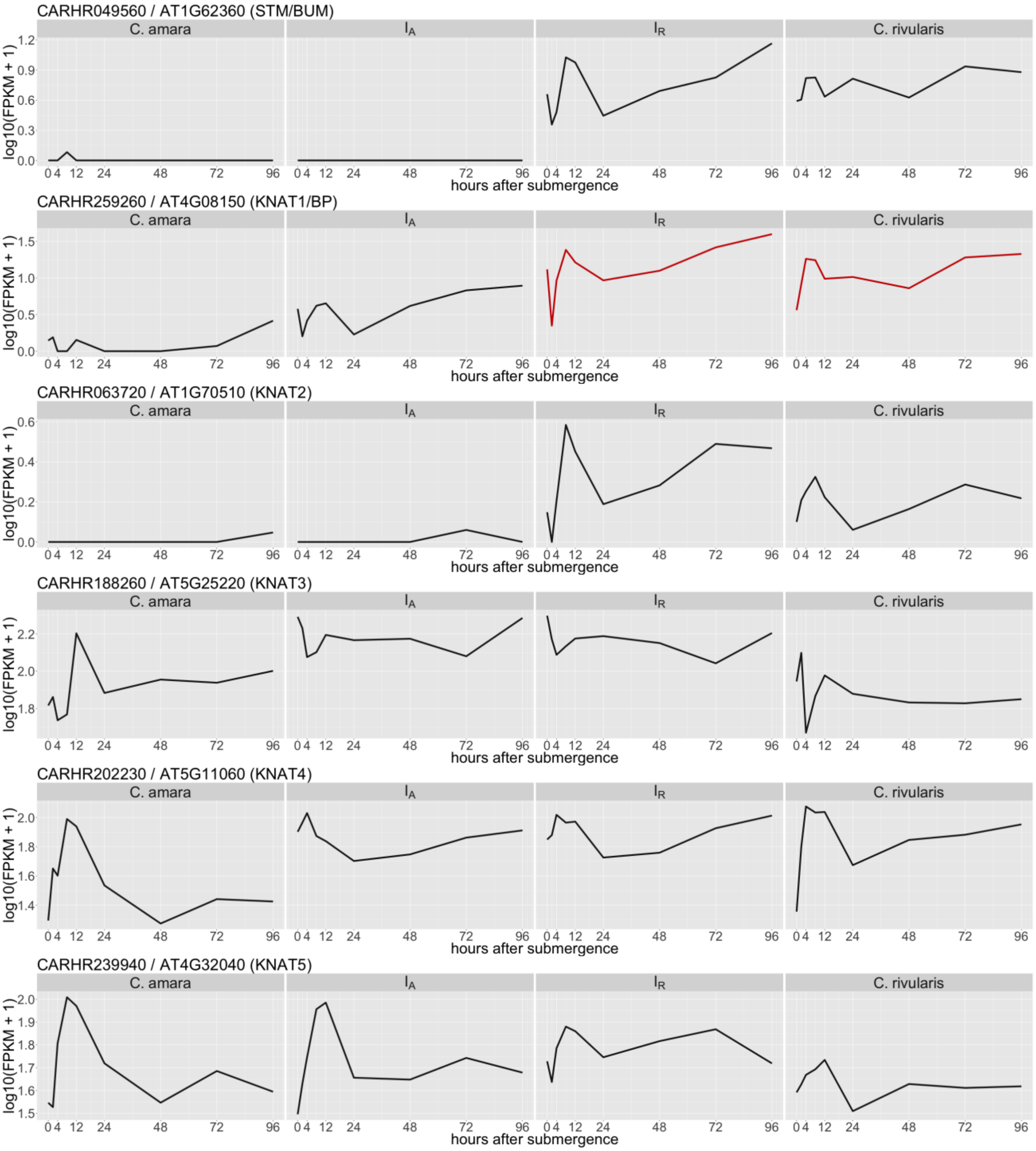

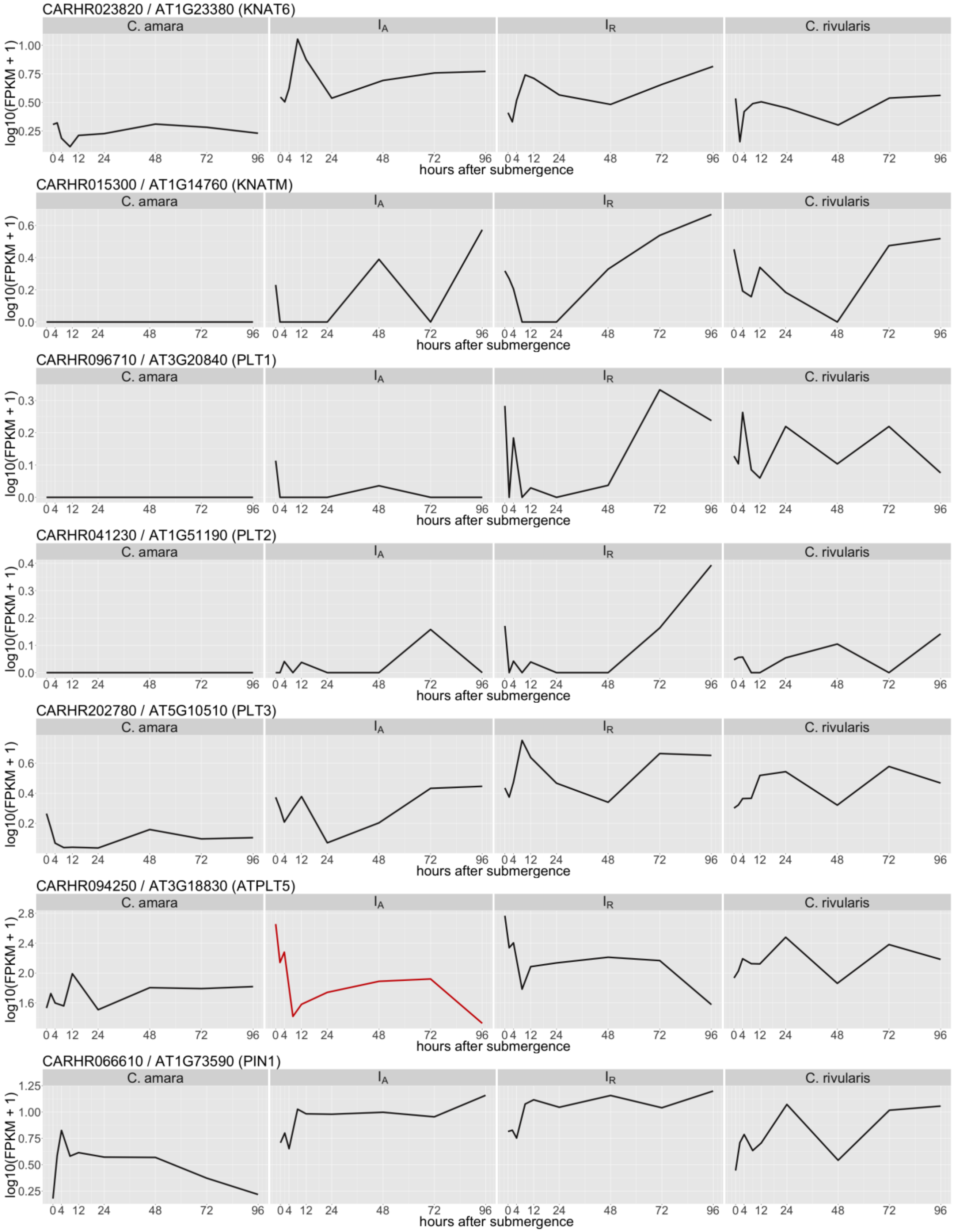
CARHR023820 / AT1G23380 (KNAT6)
CARHR015300 / AT1G14760 (KNATM)
CARHR096710 / AT3G20840 (PLT1)
CARHR041230 / AT1G51190 (PLT2)
CARHR202780 / AT5G10510 (PLT3)
CARHR094250 / AT3G18830 (ATPLT5)
CARHR066610 / AT1G73590 (PIN1)
C. amara
$I_A$
$I_R$
C. rivularis
log10(FPKM + 1)
hours after submergence
0 4 12 24 48 72 96

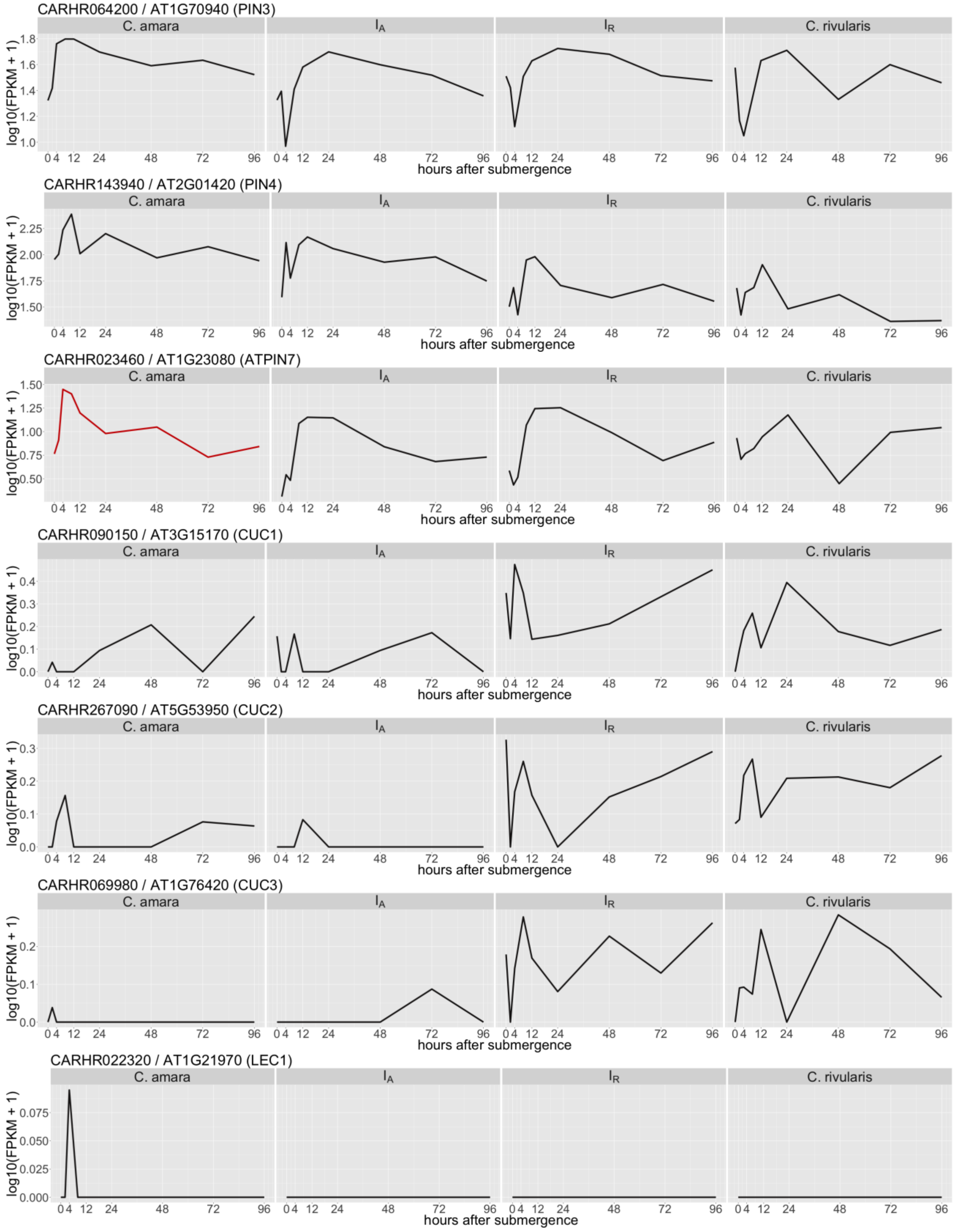

CARHR064200 / AT1G70940 (PIN3)
CARHR143940 / AT2G01420 (PIN4)
CARHR023460 / AT1G23080 (ATPIN7)
CARHR090150 / AT3G15170 (CUC1)
CARHR267090 / AT5G53950 (CUC2)
CARHR069980 / AT1G76420 (CUC3)
CARHR022320 / AT1G21970 (LEC1)
C. amara
$I_A$
$I_R$
C. rivularis
log10(FPKM + 1)
hours after submergence

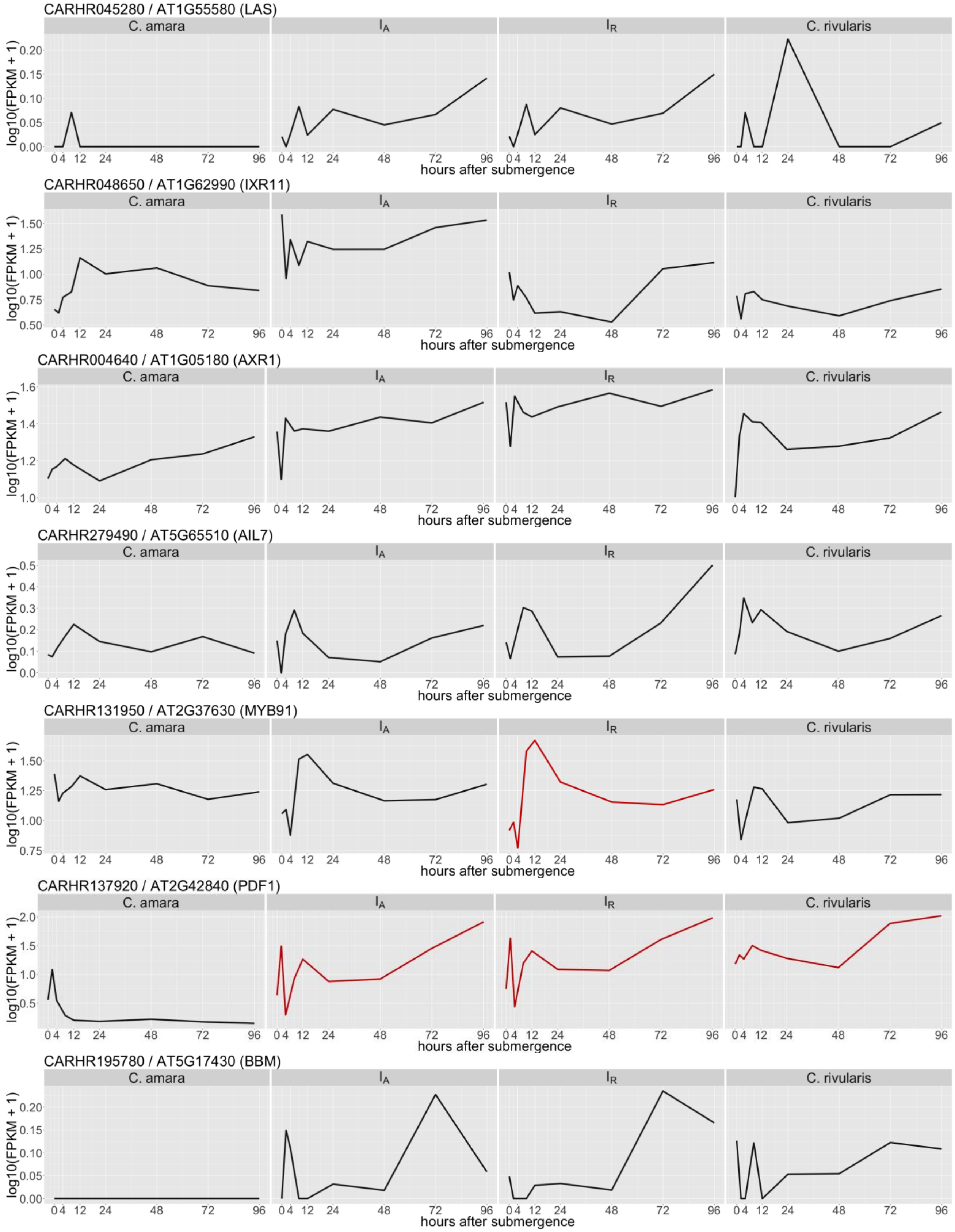
CARHR045280 / AT1G55580 (LAS)
CARHR048650 / AT1G62990 (IXR11)
CARHR004640 / AT1G05180 (AXR1)
CARHR279490 / AT5G65510 (AIL7)
CARHR131950 / AT2G37630 (MYB91)
CARHR137920 / AT2G42840 (PDF1)
CARHR195780 / AT5G17430 (BBM)
C. amara
$I_A$
$I_R$
C. rivularis
log10(FPKM + 1)
hours after submergence

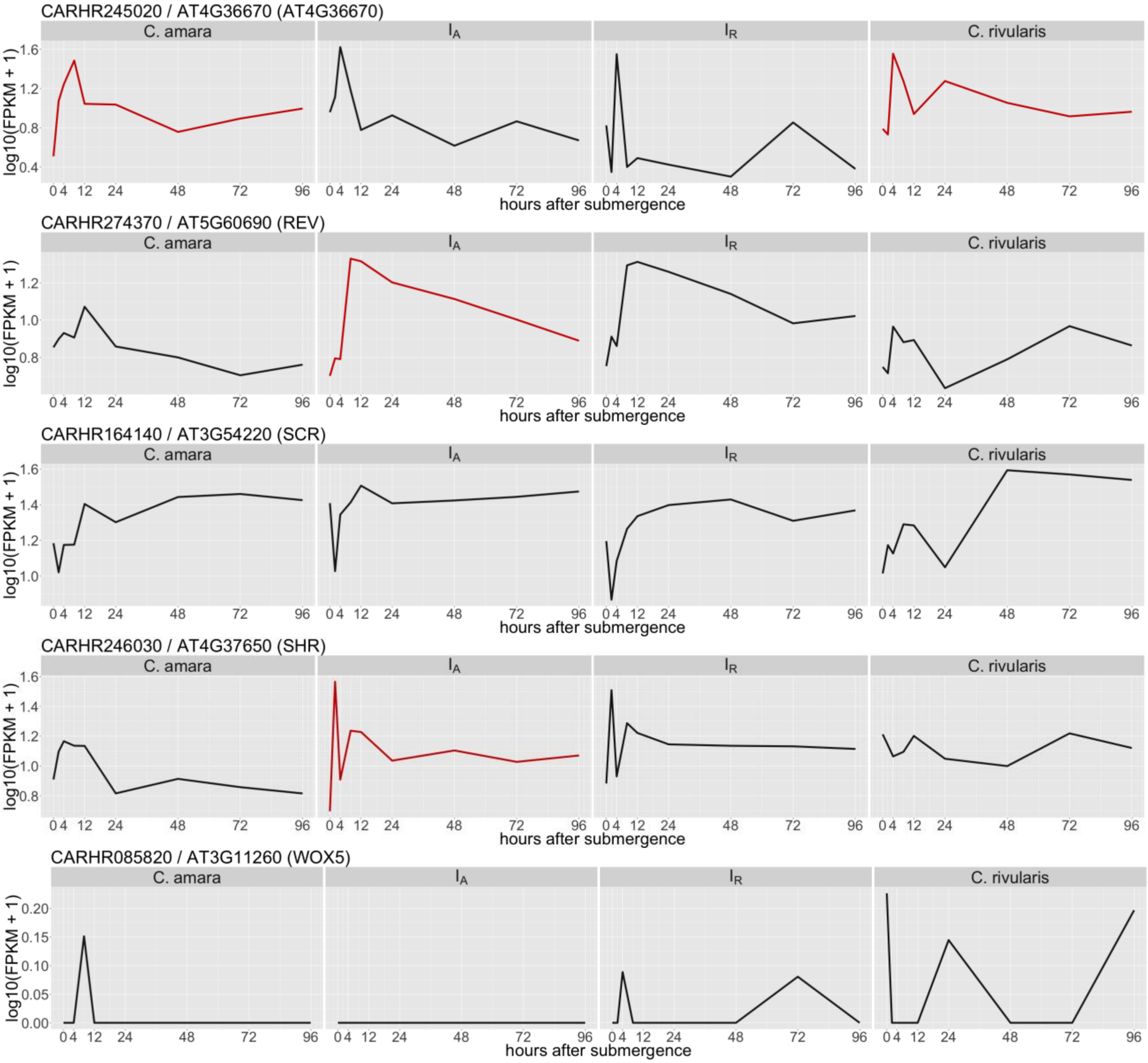
CARHR245020 / AT4G36670 (AT4G36670)
CARHR274370 / AT5G60690 (REV)
CARHR164140 / AT3G54220 (SCR)
CARHR246030 / AT4G37650 (SHR)
CARHR085820 / AT3G11260 (WOX5)
C. amara
I_A
I_R
C. rivularis
log10(FPKM + 1)
hours after submergence
0 4 12 24 48 72 96
